# A Functional Influence Based Circuit Motif That Constrains the Set of Plausible Algorithms of Cortical Function

**DOI:** 10.64898/2026.01.29.702557

**Authors:** Anna Vasilevskaya, Georg B. Keller

## Abstract

There are several plausible algorithms for cortical function that are specific enough to make testable predictions of the interactions between functionally identified cell types. Many of these algorithms are based on some variant of predictive processing. Here we set out to experimentally distinguish between two such predictive processing variants. A central point of variability between them lies in the proposed vertical communication between layer 2/3 and layer 5, which stems from the diverging assumptions about the computational role of layer 5. One assumes a hierarchically organized architecture and proposes that, within a given node of the network, layer 5 conveys unexplained bottom-up input to prediction error neurons of layer 2/3. The other proposes a non-hierarchical architecture in which internal representation neurons of layer 5 provide predictions for the local prediction error neurons of layer 2/3. We show that the functional influence of layer 2/3 cell types on layer 5 is incompatible with the hierarchical variant, while the functional influence of layer 5 cell types on prediction error neurons of layer 2/3 is incompatible with the non-hierarchical variant. Given these data, we can constrain the space of plausible algorithms of cortical function. We propose a model for cortical function based on a combination of a joint embedding predictive architecture (JEPA) and predictive processing that makes experimentally testable predictions.

## INTRODUCTION

The fact that the layered organization of neocortex is largely consistent across areas with very different function has been a key driver of the idea that all cortical areas are governed by a general algorithm (Mumford, 1992). A search for candidates for such an algorithm should be constrained to those that perform a useful computation when implemented *in silico*, generalize to all known functions of cortex, and can be mapped onto cortical cell types. The most promising proposals for such an algorithm have been different variants of predictive processing. At its core, predictive processing is based on the idea that cortex functions by learning and maintaining internal models of the world that shape predictions about the upcoming signals from the world (Mumford, 1992; Rao and Ballard, 1999). In predictive processing, these signals are typically referred to as sensory signals. We will refer to them as ***teaching signals***. The terminology of sensory signals is inadequate in non-hierarchical networks or away from the sensory periphery, where the world consists merely of other brain regions, and signals do not necessarily correspond to any specific sensory modality. Thus, following Jordan and Rumelhart (Jordan and Rumelhart, 1992) we will use the more general term of teaching signal to refer to this type of signal. We can formally define a teaching signal as a *signal that functionally serves as a reference – or a target – for a given prediction signal* (Aizenbud et al., 2025). Teaching signals are compared with their corresponding predictions, and the resulting prediction errors update the internal representation and drive plasticity to update the internal model. In its most common variant predictive processing postulates that prediction errors are computed in layer 2/3 neurons (**Figure 1A**) (Bastos et al., 2012; Keller and Mrsic-Flogel, 2018). Due to low baseline firing rates in layer 2/3, two classes of prediction error neurons were postulated – positive prediction error neurons that signal when the teaching signal exceeds the prediction, and negative prediction error neurons that signal when the prediction exceeds the teaching signal. These two classes of prediction error neurons can be mapped onto two different molecularly defined cell types of layer 2/3 (O’Toole et al., 2023). Positive and negative prediction errors exert opposing influence on internal representation neurons, a mechanism that facilitates the bidirectional integration of error signals. Internal representation neurons thus represent the estimate of the corresponding teaching signal. They are also the source of prediction and teaching signals sent to other cortical areas. Based primarily on anatomical considerations, it is often assumed that internal representation neurons are a subset of layer 5 excitatory neurons (Bastos et al., 2012; Keller and Mrsic-Flogel, 2018).

**Figure 1.**
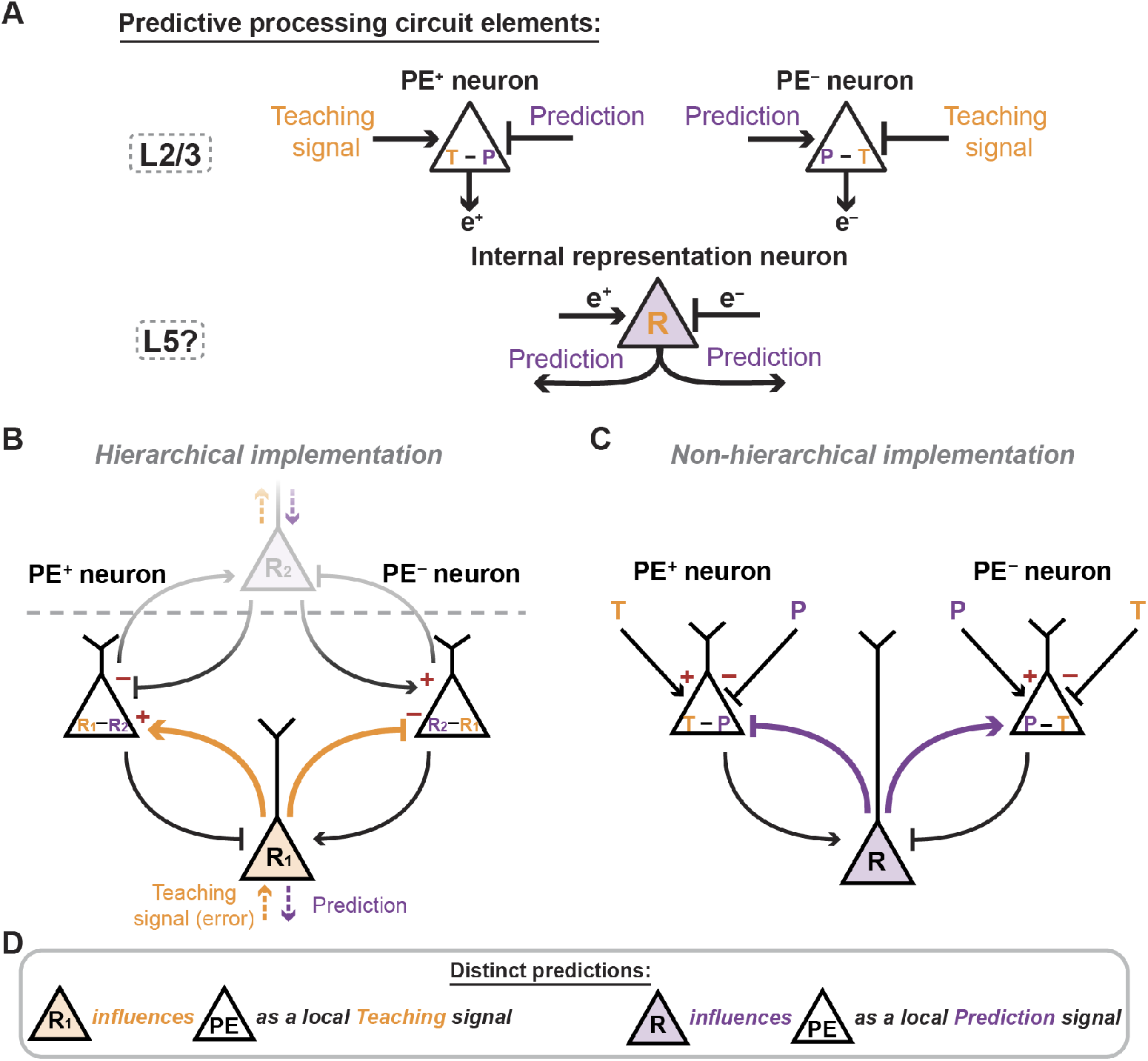
Predictive processing circuit model variants postulating a testable motif of interactions between neurons of deep and superficial cortical layers. **(A)** Two classes of neurons postulated by predictive processing. Top: A comparator circuit. Prediction error neurons compute the difference between the teaching signal and prediction. Positive prediction errors (**PE+**) are computed as Teaching signal minus Prediction (T – P), negative prediction errors (**PE-**) are computed as Prediction minus Teaching signal (P – T). Here and in the subsequent panels, arrows represent excitatory connections and blunt arrows inhibitory connections (inhibitory interneurons are omitted for simplicity – see (Attinger et al., 2017; Widmer et al., 2022) for a full circuit description). Bottom: Internal representation circuit. Internal representation neurons (**R**) integrate prediction errors and shape the prediction (or teaching) signals sent to other areas. **(B)** Schematic of the hierarchical implementation of predictive processing. **(C)** Schematic of the non-hierarchical implementation of predictive processing. **(D)** The distinct predictions made by hierarchical and non-hierarchical implementations on the functional influence of deep layer neurons on superficial layer neurons.

How prediction error neurons and representation neurons interact, however, differs in different variants of predictive processing (Spratling, 2017). The first proposal was a hierarchical variant of predictive processing in which internal representation neurons act as a source of the teaching signal for the local prediction error neurons and the next higher level of the hierarchy, while simultaneously providing a prediction signal for the preceding lower level of the hierarchy (**Figure 1B**) (Rao and Ballard, 1999). A later proposal was based on the idea of allowing for non-hierarchical arrangements of networks (Keller and Mrsic-Flogel, 2018). In this variant, the internal representation neuron interacts with the local prediction error neurons of layer 2/3 like a prediction signal (**Figure 1C**). The idea behind the non-hierarchical implementation was two-fold. First, each node of the network should be able to work independently. With the internal representation acting as a local prediction, a network with just one node will function like a simple temporal difference detector. Second, the exchange of teaching signals and predictions should be possible between any two nodes in a network. Predictions exchanged between primary sensory areas (Garner and Keller, 2022), are likely an example of a non-hierarchical interaction where predictions and teaching signals are exchanged bidirectionally. The non-hierarchical variant of predictive processing anticipates that error neurons can integrate over multiple sources of predictions. It should be noted that neither model, of course, represents the only possible solution for hierarchical/non-hierarchical organization of cortical nodes, we just use this terminology to name the two models based on their most important conceptual difference. As a result of the differences in implementation, the two models make experimentally distinguishable predictions about the interactions between representation neurons and prediction error neurons. Essential is the fact that all proposed interactions between prediction error neurons and local internal representation neurons have reversed sign in the two models (**Figures 1B and 1C**).

In this work we focus on testing these predictions in mouse primary visual cortex (V1) in the context of visuomotor interactions. Our approach was based on the ability to functionally and molecularly identify layer 2/3 positive and negative prediction error neurons and probe their interaction with other genetically identified neuron types in cortex using optogenetics. If the hierarchical model is correct, we expect to find a genetically identified neuron type (the still unidentified internal representation neuron) in the local circuit that excites positive prediction error neurons and inhibits negative prediction error neurons, while at the same time we expect this representation neuron type to be inhibited by positive prediction error neurons and excited by negative prediction error neurons (**Figure 1B**). If the non-hierarchical model is correct, we expect to find a genetically identified neuron type in the local circuit that inhibits positive prediction error neurons and excites negative prediction error neurons, again with reversed influence in the opposite direction (**Figure 1C**).

## RESULTS

In an initial set of experiments, we aimed to test the predictions made by the two circuit models (**Figure 1**) regarding the functional influence that internal representation neurons exert on local prediction error neurons. Critical for the interpretation of our results here is the assumption that internal representation neurons are a genetically identifiable cell type in layers 5 or 6. Thus, we screened the functional influence of genetically identified subsets of excitatory neurons in layers 5 and 6 on functionally identified prediction error neurons in layer 2/3. To be able to record the activity of layer 2/3 neurons, we expressed a genetically encoded calcium indicator by injecting an AAV2/1-EF1α-GCaMP6f vector in visual cortex. We did this in three different mouse lines that express Cre recombinase in a subset of layer 5 intratelencephalic (IT, Tlx3-Cre), layer 5 extratelencephalic (ET, Fezf2-Cre) or layer 6 (Ntsr1-Cre) neurons. To be able to optogenetically activate these neurons, we also injected an AAV2/1-hSyn-DIO-ChrimsonR-tdTomato vector to express the optogenetic activator ChrimsonR in Cre positive neurons in primary visual cortex (**Figure 2A**). For all experiments, mice were head fixed on a spherical treadmill surrounded by a toroidal screen that displayed a virtual reality environment. The virtual environment consisted of a tunnel, with walls patterned with vertical sinusoidal gratings (**Figure 2B**). We used a two-photon microscope with a coaxial 637 nm stimulation laser to activate ChrimsonR while recording calcium responses in layer 2/3 neurons (**Methods**).

**Figure 2.**
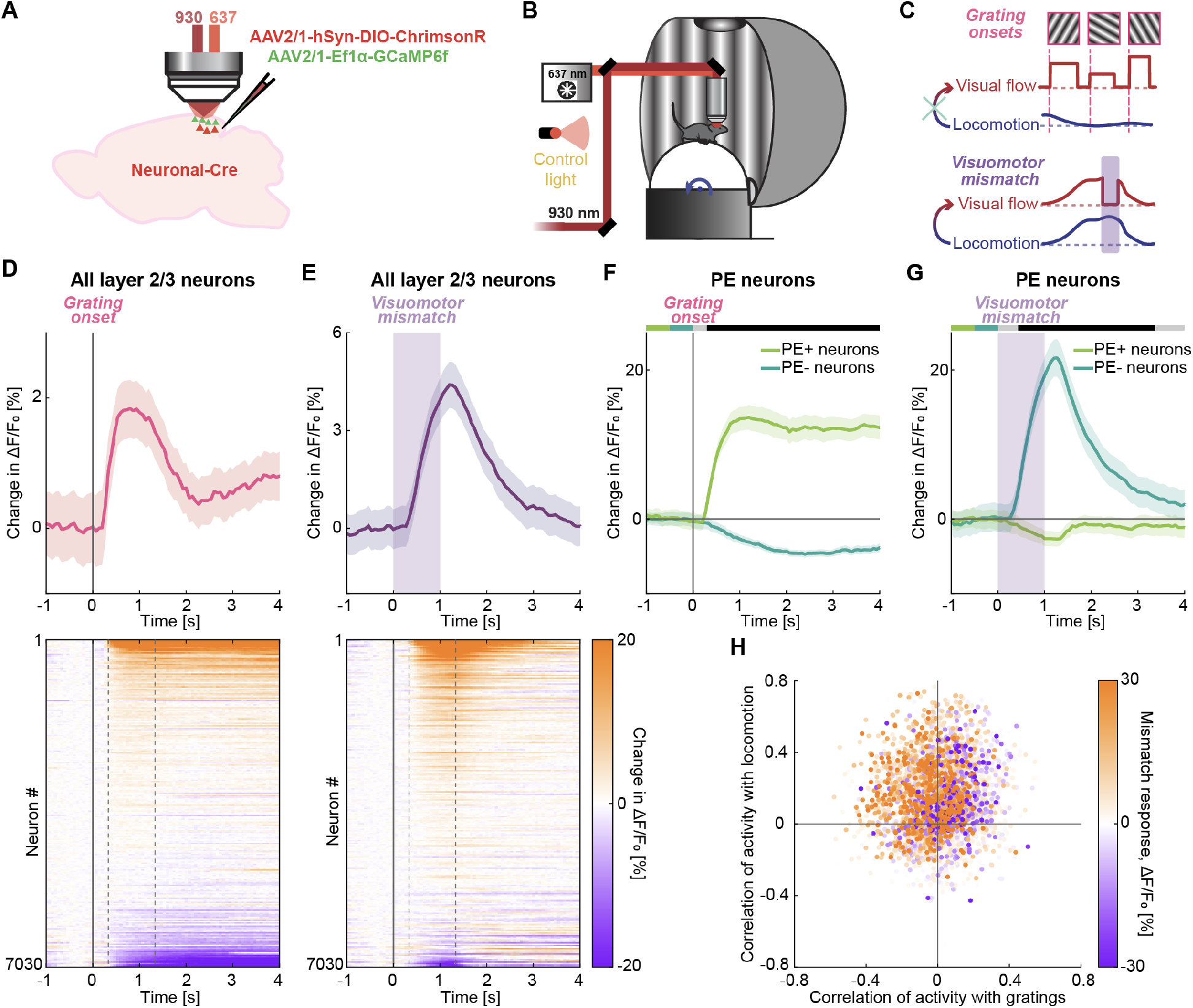
Functional identification of positive and negative prediction error neurons in layer 2/3 of V1. **(A)** AAV vector injections were used to express ChrimsonR-tdTomato in different Cre-positive deep layer neurons (**Figures 3-5**), and GCaMP6f in layer 2/3 neurons of V1. The ChrimsonR virus was omitted in no-ChrimsonR control mice. **(B)** Schematic of the two-photon microscope and a virtual reality system. **(C)** Schematic of the visuomotor stimuli used to probe for prediction error responses. Grating onsets were used to identify positive prediction error neurons and visuomotor mismatch stimuli were used to identify negative prediction error neurons. **(D)** Top: Mean layer 2/3 population response to grating onset. Here and in the subsequent panels, the solid line represents the hierarchical bootstrap estimates of the mean trace. Shading around the mean is the bootstrap error, defined as one standard deviation of the bootstrap distribution at each time bin. Bottom: Heatmap of mean responses of all layer 2/3 neurons. Neurons are sorted by the amplitude of their grating onset response, dashed gray lines mark the sorting window. **(E)** As in **D**, but for visuomotor mismatch. Purple shading marks the duration of the mismatch stimulus. **(F)** Mean response of positive (light green) and negative (dark green) prediction error neurons to grating onset. Here and in the subsequent panels, response curves are compared for each time bin: in the horizontal bars above the plot, black marks time bins where p<0.05 and gray mark time bins where p>0.05. Colored bars to the left indicate which lines are compared. **(G)** Mean response of positive (light green) and negative (dark green) prediction error neurons to visuomotor mismatch. **(H)** Scatter plot of the correlation between neuronal activity and optical flow speed during grating sessions against the correlation between neuronal activity and locomotion speed for all layer 2/3 neurons. Dot color indicates the amplitude of the response to visuomotor mismatch.

To functionally identify prediction error neurons in layer 2/3 of V1, we used the virtual reality system to present a set of visuomotor stimuli that are known to elicit different types of prediction error responses in mouse V1 (Jordan and Keller, 2020; O’Toole et al., 2023). Mice were exposed first to a closed loop condition, in which visual flow feedback in the virtual tunnel was coupled to locomotion on a spherical treadmill. In the closed loop condition, we presented brief (1 s) unpredictable halts in visual flow, thus briefly breaking the coupling between locomotion and visual flow feedback. We refer to such a break in coupling as a visuomotor mismatch. The responses to a visuomotor mismatch can be interpreted as signaling a negative prediction error (Keller and Mrsic-Flogel, 2018; Keller and Sterzer, 2024; Rao and Ballard, 1999), since during a visuomotor mismatch there is less visual flow than predicted given locomotion (**Figure 2C**, bottom). After the closed loop condition, mice were exposed to an open loop condition. In the open loop condition, locomotion and visual flow feedback were uncoupled and mice viewed a replay of the visual flow that they had self-generated in the preceding closed loop session. Finally, during a grating session, full-field gratings of randomized orientation, direction, and duration were presented to the mouse at random times (**Methods**). In both open loop and grating sessions, the visual stimuli were presented independently of the locomotion behavior of the mouse.

The presentation of unpredictable grating stimuli results in increases of activity in a subset of layer 2/3 neurons that are typically referred to as visual, or sensory, responses. As these stimuli constitute more visual input than predicted by movement (**Figure 2C**, top), such responses can be interpreted as positive prediction error responses (Keller and Mrsic-Flogel, 2018; Keller and Sterzer, 2024; Rao and Ballard, 1999). Note, in predictive processing multiple neuron types are expected to have sensory driven increases in activity, positive prediction error neurons are just one of these (Keller and Mrsic-Flogel, 2018). As a note on terminology, we will refer to ‘visuomotor mismatch’ and ‘grating onset’ responses, after the stimuli used to trigger them, and to ‘negative and positive prediction error neurons’, when talking about the neurons in layer 2/3 that are responsive to these stimuli.

Both visuomotor mismatch and grating onset elicited strong population responses in layer 2/3. For each stimulus, the distribution of individual neuronal responses was composed of a continuum, ranging from strong increases in activity to decreases in activity (**Figures 2D and 2E**). We selected the 15% of neurons most responsive to grating onset and will refer to these as positive prediction error neurons. By construction, positive prediction error neurons exhibited larger grating onset responses than the population average (**Figures 2D and 2F**) but exhibited negative responses to visuomotor mismatch (**Figure 2G**). Similarly, we selected the 15% most responsive neurons to visuomotor mismatch and will refer to these as negative prediction error neurons. Note, in both cases our results do not depend on the exact percentage of neurons included here (**Figures S3, S6, S7, and S9**). Consistent with previous findings (Attinger et al., 2017), negative prediction error neurons exhibited a reduction in response to the presentation of full-field gratings (**Figure 2F**). Also as previously reported (Widmer et al., 2022), their responses to locomotion onset depended on whether the onset occurred in closed or open loop conditions (**Figure S1**). Locomotion onset responses of layer 2/3 were on average stronger in the open loop condition (including grating sessions) than in the closed loop condition (**Figure S1A**), possibly through a visual flow induced inhibition – since a closed loop locomotion onset is always accompanied by a visual flow onset. Locomotion onset responses of positive prediction error neurons were smaller than those of negative prediction error neurons and were stronger in closed loop than in open loop (**Figure S1B**). This response profile is also consistent with the functional and computational features of positive prediction error neurons that are thought to compute the difference between visual teaching input and motor related prediction (Jordan and Keller, 2020; Keller and Mrsic-Flogel, 2018). Finally, prediction error neurons in layer 2/3 exhibit differential correlations with visual flow and locomotion speed in open loop conditions (Attinger et al., 2017). Plotting the distribution of layer 2/3 neuronal activity correlations with a binarized grating trace against the activity correlations with locomotion speed shows that mismatch responsive neurons preferentially exhibit negative correlations with visual input and positive correlations with locomotion (**Figure 2H**). An opposing influence of a negative visual teaching input and a positive motor related prediction signal is a hallmark feature of negative prediction error neurons in V1 (Jordan and Keller, 2020).

Thus, we can separate layer 2/3 neurons into putative positive and negative prediction error neurons based on their functional responses. To test the functional influence of layer 5 IT neurons on these subpopulations of layer 2/3 neurons, we stimulated the Tlx3 neurons optogenetically at random times while recording layer 2/3 responses (**Figure 3A**). We found that we could elicit responses in layer 2/3 neurons when stimulating layer 5 IT neurons (**Figure 3B**). To control for responses that were elicited independently of the optogenetic activation, we used both a control light outside of the craniotomy, to emulate purely visual responses in the same mice, and a ‘no-ChrimsonR’ control using the stimulation light in a different cohort of mice, in which we omitted the injection of the ChrimsonR vector. In both control conditions, we found visual responses to the control or the stimulation light turning on (**Figure 3B**). Thus, even though the stimulation light is red (637 nm), and outside the peak sensitivity of the mouse retina, it appears to be bright enough to be visible to the mice. However, these responses were considerably smaller than the optogenetic stimulation responses (**Figure 3B**) and activated a different set of neurons. Sorting neurons by their response strength to optogenetic stimulation of layer 5 IT neurons revealed a spectrum of responses (**Figure 3C**). Responses of the same neurons to control light stimulation were only poorly correlated with the responses to layer 5 IT stimulation (**Figures 3C and S2A**). In the no-ChrimsonR control mice, the response distribution was similar to that in the control light stimulation case (**Figure S2B**). Based on this, we conclude that the optogenetic activation of layer 5 IT neurons triggers responses in layer 2/3 neurons. Note, while there are several indirect pathways from layer 5 IT to layer 2/3 that could contribute to this influence, we see no reason to assume that the majority of the influence is not via direct projections from layer 5 to layer 2/3 (Hage et al., 2022; MICrONS Consortium, 2025).

**Figure 3.**
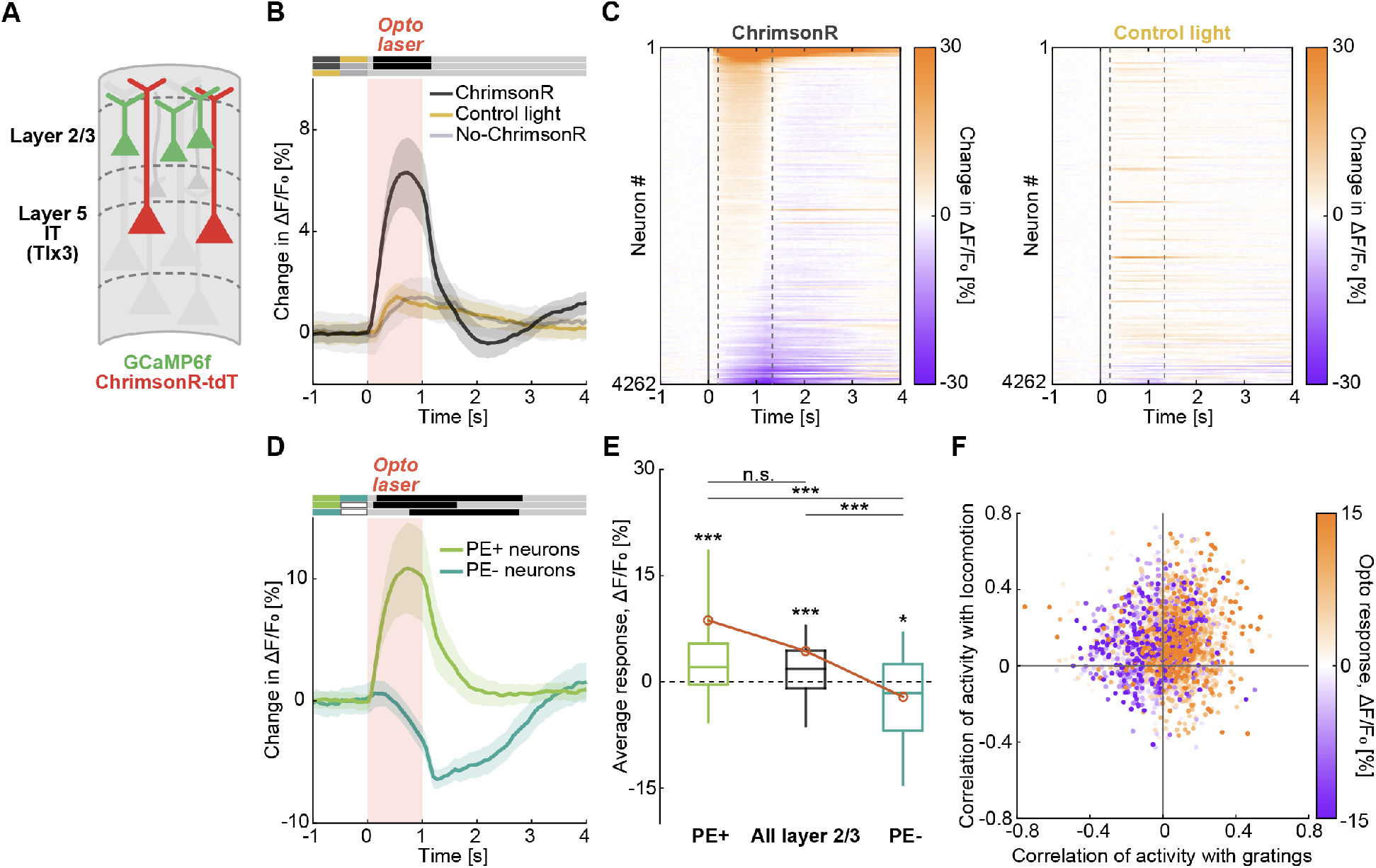
Opposing functional influence of Tlx3 layer 5 neurons on positive and negative prediction error neurons of layer 2/3. **(A)** We optogenetically activated Tlx3 layer 5 neurons while recording calcium activity in layer 2/3 neurons. **(B)** Mean layer 2/3 population responses to optogenetic stimulation of Tlx3 layer 5 neurons (dark gray), control light stimulation in the same mice (yellow), and stimulation in no-ChrimsonR control mice (light gray). Red shading marks the stimulation interval. **(C)** Heatmap of mean layer 2/3 neuronal responses to optogenetic stimulation of Tlx3 layer 5 neurons (left) and control light stimulation (right). Neurons are sorted by the amplitude of their response to the optogenetic stimulation. **(D)** Mean responses of positive (light green) and negative (dark green) prediction error neurons in layer 2/3 to optogenetic stimulation of Tlx3 layer 5 neurons. Red shading marks the optogenetic stimulation interval. **(E)** Box plots showing average responses of positive prediction error neurons (light green), negative prediction error neurons (dark green), and all layer 2/3 neurons (black) to optogenetic stimulation of Tlx3 layer 5 neurons. Boxes indicate the interquartile range, the central line marks the median, and whiskers the 10th and 90th percentiles. Orange circles mark the mean response. Here and elsewhere, n.s.: not significant, *: p<0.05, **: p<0.01, ***: p<0.001. **(F)** Scatter plot of the correlation between neuronal activity and optical flow speed against the correlation between neuronal activity and locomotion speed during grating sessions for all layer 2/3 neurons. Dot color indicates the amplitude of the response to optogenetic stimulation of Tlx3 layer 5 neurons.

To test predictions of the two circuit model types (**Figure 1**) and determine whether the functional influence of layer 5 on layer 2/3 depends on the functional response profile of the layer 2/3 neuron, we analyzed the stimulation responses of positive and negative prediction error neurons (as defined in **Figure 2**) separately. We found that positive prediction error neurons increased their activity on average, while negative prediction error neurons decreased their activity in response to layer 5 IT stimulation (**Figures 3D and 3E**). Again, we plotted the correlation of activity with gratings against that with locomotion for each neuron, and color coded the response strength to optogenetic stimulation of layer 5 IT neurons (**Figure 3F**). Similar to the pattern of mismatch responses, we found that neurons that decrease activity on stimulation of layer 5 IT neurons preferentially exhibited positive correlation with locomotion and negative correlation with gratings. This functionally specific modulation of layer 2/3 neurons was preserved if we used different fractions of mismatch and grating onset responsive neurons (**Figures S3A-S3C**). In contrast, no such modulation was observed with control light stimulation (**Figure S3D**) or in no-ChrimsonR control mice (**Figure S4**). Assuming layer 5 IT neurons are internal representation neurons, this would be exactly the relationship postulated by the hierarchical variant of predictive processing (**Figure 1B**), and the opposite of influence postulated by the non-hierarchical variant (**Figure 1C**). Thus, we can conclude that the influence of layer 5 IT neurons on layer 2/3 is consistent with the hierarchical variant of predictive processing under the assumption that layer 5 IT neurons are internal representation neurons, but inconsistent with the non-hierarchical variant.

The stimuli we used to identify positive and negative prediction error neurons, visuomotor mismatch and grating onset, only activate a subset of all prediction error neurons. Both sensory and visuomotor mismatch responses (Zmarz and Keller, 2016) exhibit tuning, such that an individual neuron will only respond to a subset of all possible stimuli. While it is not experimentally feasible to cover the full stimulus space, we began to address whether the opposing influence of layer 5 IT stimulation generalizes to other visual and mismatch stimuli. To this end, we repeated the analysis using a limited alternative set of visual and mismatch stimuli. In the case of visual stimuli, we can take advantage of the fact that layer 2/3 neurons are tuned to grating orientation (**Figure S5A**). Independent of which grating orientation we use to identify positive prediction error neurons, we find the same opposing influence of layer 5 IT stimulation (**Figure S5B**). For visuomotor mismatch responses, we performed an additional set of experiments in which we exposed mice both to normal coupling, with forward locomotion resulting in backward visual flow, and to an inverted coupling, where forward locomotion resulted in forward visual flow (**Figure S5C**). In both conditions we measured visuomotor mismatch responses in layer 2/3 neurons. We found that the two types of visuomotor mismatch tended to activate different sets of neurons (**Figure S5D**). Like normal visuomotor mismatch neurons, inverted visuomotor mismatch neurons also exhibited a decrease in activity upon visual stimulation (**Figure S5E**). Importantly, we found that inverted visuomotor mismatch neurons also exhibited the same sign of layer 5 IT influence we had seen for normal visuomotor mismatch neurons (**Figure S5F**). Thus, we speculate that layer 5 IT neurons generally have an excitatory functional influence on positive prediction error neurons and an inhibitory functional influence on negative prediction error neurons in layer 2/3.

Our interpretation of the results relies on an assumption of which cell type functions as internal representation neuron. It is possible that a different cell type in layers 5 or 6 might exert an inverted pattern of influence on layer 2/3. Thus, we repeated our experiments with a Cre line that labels layer 5 ET neurons (Fezf2-Cre) and one that labels layer 6 neurons (Ntsr1-Cre). Interestingly, we found the same opposing influence of these neuron types on positive and negative prediction error neurons in layer 2/3. Positive prediction error neurons increased their activity upon stimulation of both layer 5 ET (**Figures 4 and S6**) and layer 6 neurons (**Figures 5 and S7**), while negative prediction error neurons decreased their activity on stimulation – very similar to what we found for layer 5 IT neurons (**Figure 3**). Thus, we found no evidence of an interaction between deep cortical layers and layer 2/3 consistent with internal representation neurons and the non-hierarchical variant of predictive processing (**Figure 1C**), but rather all interactions are consistent with internal representation neurons and the hierarchical variant of predictive processing (**Figure 1B**). Hence, quite remarkably, we find that the functional response profile of layer 2/3 neurons is predictive of the influence that they receive from stimulation of deep cortical layers. Moreover, this influence is consistent with that predicted by the hierarchical predictive processing model.

**Figure 4.**
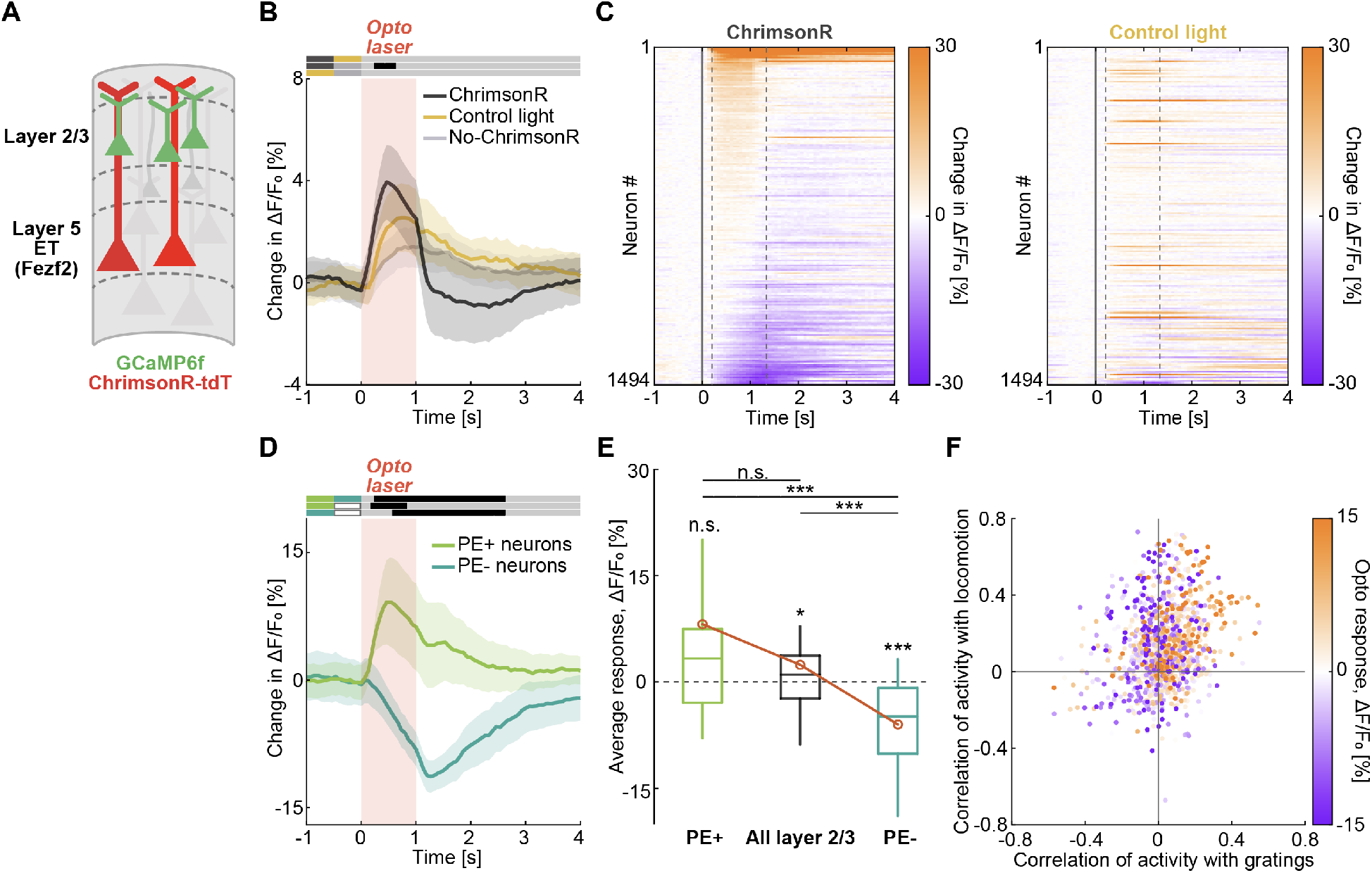
Opposing functional influence of Fezf2 layer 5 neurons on positive and negative prediction error neurons of layer 2/3. **(A)** We optogenetically activated Fezf2 layer 5 neurons while recording calcium activity in layer 2/3 neurons. **(B)** Mean layer 2/3 population responses to optogenetic stimulation of Fezf2 layer 5 neurons (dark gray), control light stimulation in the same mice (yellow), and stimulation in no-ChrimsonR control mice (light gray). Red shading marks the stimulation interval. **(C)** Heatmap of mean layer 2/3 neuronal responses to optogenetic stimulation of Fezf2 layer 5 neurons (left) and control light stimulation (right). Neurons are sorted by the amplitude of their response to the optogenetic stimulation. **(D)** Mean responses of positive (light green) and negative (dark green) prediction error neurons in layer 2/3 to optogenetic stimulation of Fezf2 layer 5 neurons. Red shading marks the optogenetic stimulation interval. **(E)** Box plots showing average responses of positive prediction error neurons (light green), negative prediction error neurons (dark green), and all layer 2/3 neurons (black) to optogenetic stimulation of Fezf2 layer 5 neurons. Boxes indicate the interquartile range, the central line marks the median, and whiskers the 10th and 90th percentiles. Orange circles mark the mean response. **(F)** Scatter plot of the correlation between neuronal activity and optical flow speed against the correlation between neuronal activity and locomotion speed during grating sessions for all layer 2/3 neurons. Dot color indicates the amplitude of the response to optogenetic stimulation of Fezf2 layer 5 neurons.

**Figure 5.**
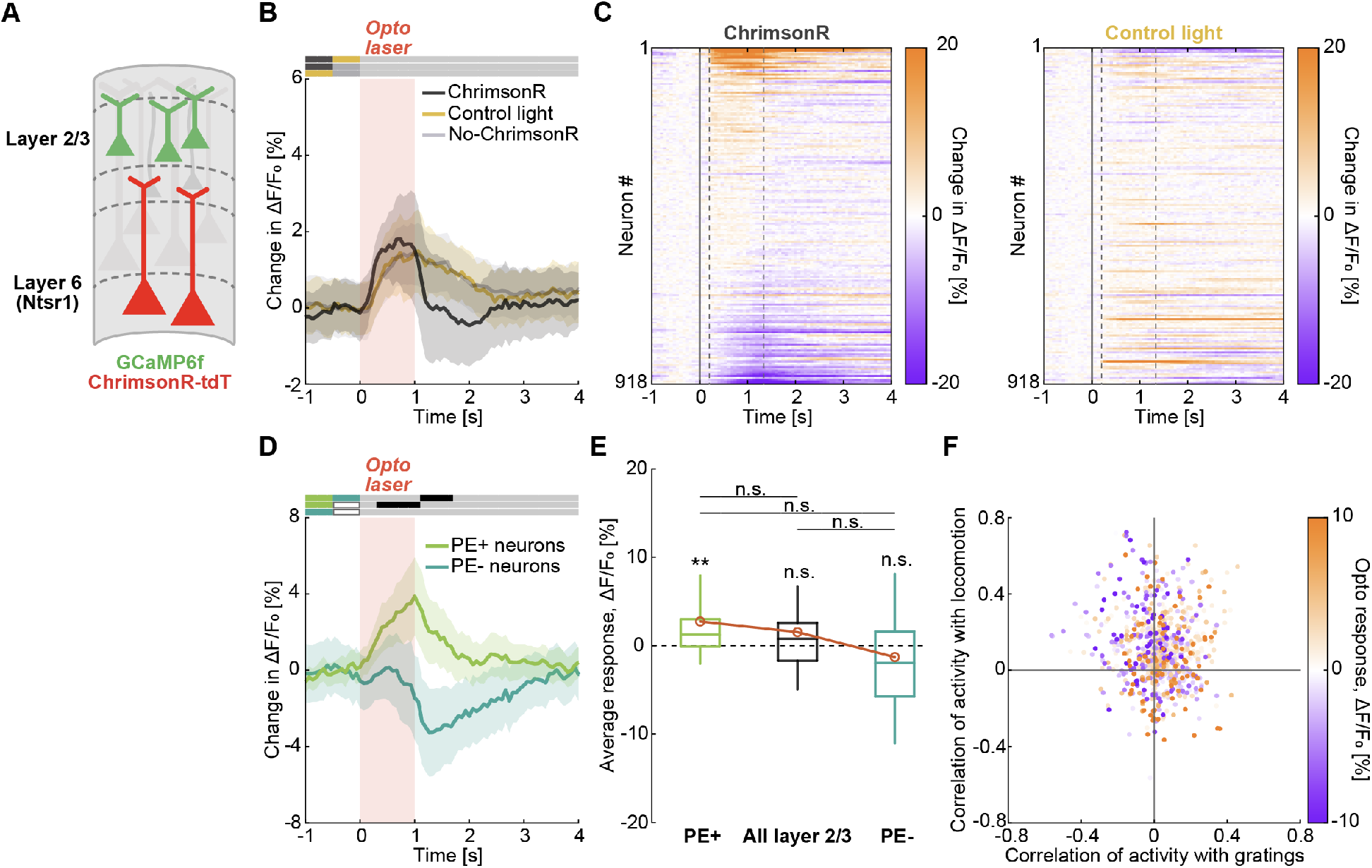
Functional influence of Ntsr1 layer 6 neurons on positive and negative prediction error neurons of layer 2/3. **(A)** We optogenetically activated Ntsr1 layer 6 neurons while recording calcium activity in layer 2/3 neurons. **(B)** Mean layer 2/3 population responses to optogenetic stimulation of Ntsr1 layer 6 neurons (dark gray), control light stimulation in the same mice (yellow), and stimulation in no-ChrimsonR control mice (light gray). Red shading marks the stimulation interval. **(C)** Heatmap of mean layer 2/3 neuronal responses to optogenetic stimulation of Ntsr1 layer 6 neurons (left) and control light stimulation (right). Neurons are sorted by the amplitude of their response to the optogenetic stimulation. **(D)** Mean responses of positive (light green) and negative (dark green) prediction error neurons in layer 2/3 to optogenetic stimulation of Ntsr1 layer 6 neurons. Red shading marks the optogenetic stimulation interval. **(E)** Box plots showing average responses of positive prediction error neurons (light green), negative prediction error neurons (dark green), and all layer 2/3 neurons (black) to optogenetic stimulation of Ntsr1 layer 6 neurons. Boxes indicate the interquartile range, the central line marks the median, and whiskers the 10th and 90th percentiles. Orange circles mark the mean response. **(F)** Scatter plot of the correlation between neuronal activity and optical flow speed against the correlation between neuronal activity and locomotion speed during grating sessions for all layer 2/3 neurons. Dot color indicates the amplitude of the response to optogenetic stimulation of Ntsr1 layer 6 neurons.

To further test these models, we investigated the predictions they make about the influence in the reverse direction, from prediction error neurons on internal representation neurons (**Figure 6A**). This is technically a bit more challenging as it requires a way to target optogenetic stimulation specifically either to positive or to negative prediction error neurons in layer 2/3. To do this, we used two different artificial regulatory elements that can be used in AAV vectors to bias expression of a reporter to either positive or negative prediction error neurons (O’Toole et al., 2023) (**Figure 6B**). The AP.Baz1a.1 regulatory element biases expression to positive prediction error neurons (the Rrad transcriptional cell type), and the AP.Adamts2.1 regulatory element biases expression to negative prediction error neurons (the Adamts2 transcriptional cell type). We will refer to neurons labeled using this strategy by the target cell type of the corresponding artificial regulatory element (Rrad and Adamts2). We performed these experiments in two separate cohorts of mice. In the first, we expressed ChrimsonR under the control of AP.Baz1a.1 regulatory element and recorded calcium responses in Tlx3 layer 5 IT neurons (**Figure 6C**). We found that optogenetic activation of Rrad neurons resulted in a trend towards increased activity in layer 5 IT neurons (**Figures 6D and 6G**). Repeating the experiments in mice that expressed ChrimsonR under the control of the AP.Adamts2.1 regulatory element (**Figure 6E**), we found that optogenetic activation of Adamts2 neurons resulted in a net decrease of activity in layer 5 IT neurons (**Figures 6F and 6G**). This interaction is the inverse of what the hierarchical version of predictive processing would postulate under the assumption that layer 5 neurons function as internal representation neurons, but matches the predictions made by the non-hierarchical variant under the same assumption (**Figure 6A**).

**Figure 6.**
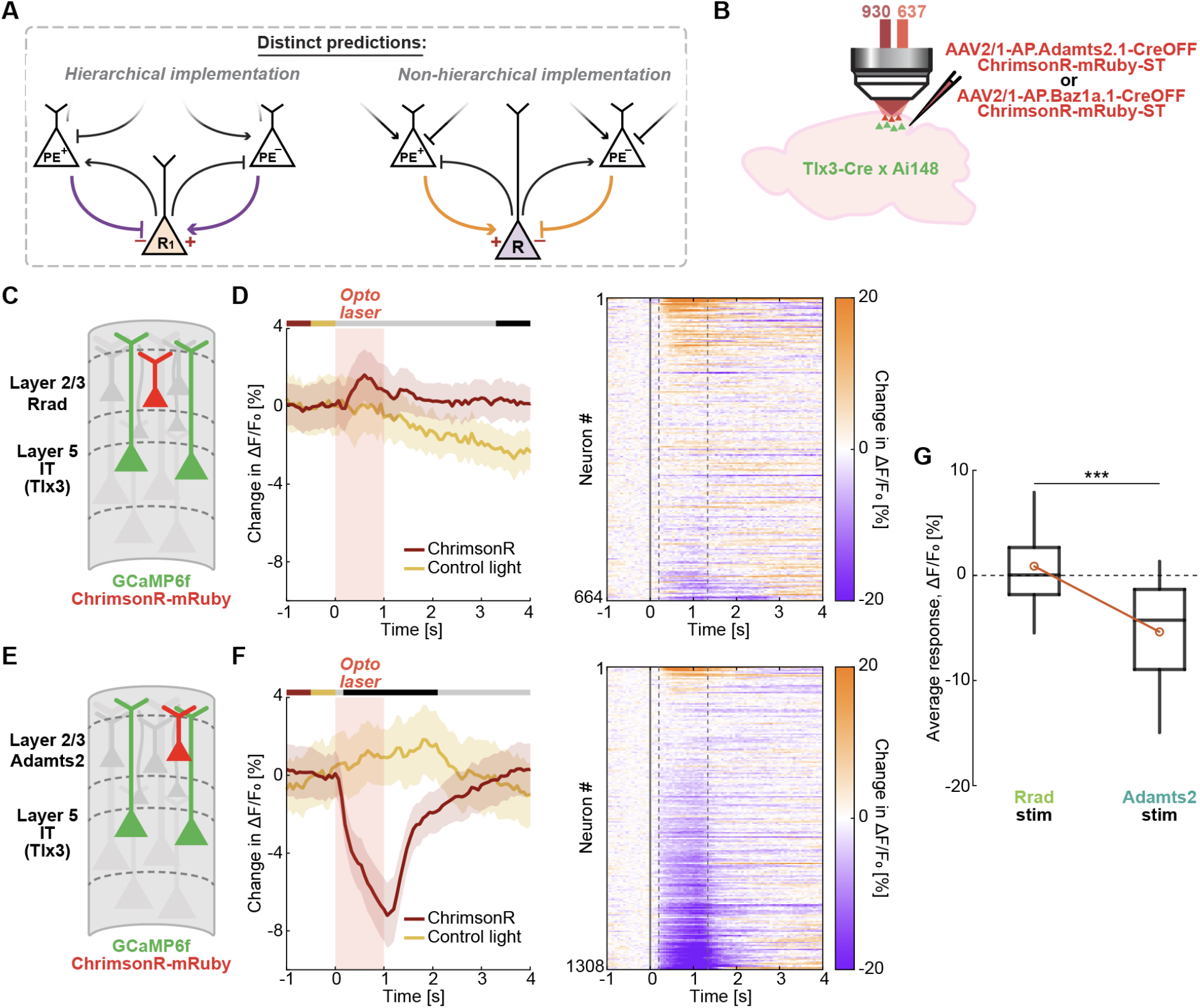
Opposing functional influence of positive and negative prediction error neurons on Tlx3 layer 5 IT neurons. **(A)** Schematic of the distinct experimental predictions made by hierarchical and non-hierarchical predictive processing implementations on the functional influence of superficial layer neurons on deep layer neurons. **(B)** Two artificial regulatory elements (either AP.Baz1a.1 or AP.Adamts2.1) were used in AAV vectors to bias expression of soma-targeted (ST) ChrimsonR to either positive (Rrad) or negative (Adamts2) prediction error neurons respectively. AAV vectors were injected in V1 of Tlx3-Cre x Ai148 mice. **(C)** We optogenetically activated Rrad neurons while recording calcium activity in Tlx3 layer 5 IT neurons. **(D)** Left: Mean layer 5 IT populational response to optogenetic stimulation of Rrad neurons (red) and control light stimulation in the same mice (yellow). Red shading marks the stimulation interval. Right: Heatmap of mean layer 5 IT neuronal responses to optogenetic stimulation of Rrad neurons. Neurons are sorted by the amplitude of their response to the optogenetic stimulation. **(E)** We optogenetically activated Adamts2 layer 2/3 neurons while recording calcium activity in Tlx3 layer 5 IT neurons. **(F)** As in **D**, but for stimulation of Adamts2 neurons. **(G)** Comparison of the responses of Tlx3 layer 5 IT neurons to optogenetic stimulation of Rrad or Adamts2 neurons. Boxes indicate the interquartile range, the central line the median, and whiskers the 10th and 90th percentiles. Orange circles mark the mean response.

Where does that leave us? Neither variant of the model can account for the influence between superficial and deep layers under the assumption that prediction error neurons are in layer 2/3 and internal representation neurons in layers 5 or 6. There are a number of caveats to this conclusion that originate in the fact that our approach may not have covered all possible subtypes of deep cortical neurons (see **Discussion**). But let us call this a lemma for now: either both predictive processing models are wrong, or internal representation neurons are not in deep cortical layers.

As a starting point for our argument to explain these observations, it is important to note that from the perspective of prediction error neurons in layer 2/3, the influence of deep cortical layers is consistent with a teaching signal. Based on this, we speculated that the prediction error responses in layer 2/3 may not be computed with respect to sensory input, but with respect to layer 5 activity as a teaching signal. We noticed that the functional influence of layer 5 IT neurons depended on whether the mice were locomoting. The functional influence was decreased while mice were locomoting, but in such a way that there was a population of neurons that decreased their activity stronger during locomotion than during stationary periods (**Figures S8A-S8C**). These differences could not be explained by differences in the optogenetic activation of the layer 5 IT neurons themselves, since they were similarly activated in both cases (**Figures S8D-S8F**). Such an unmasking of an inhibitory functional influence would be consistent with a computation in the layer 2/3 negative prediction error neurons that compare an excitatory prediction with an inhibitory teaching signal from layer 5.

Given our results thus far, we hypothesized that the predictive processing network in superficial layers of cortex would function to model and predict activity in deep cortical layers. One way to investigate this idea more directly is to generate artificial activity patterns in layer 5 and test whether layer 2/3 responses look like prediction errors with respect to perturbations of the artificial layer 5 activity patterns. Thus, we designed an experiment in which the optogenetic activation of layer 5 IT neurons was closed loop coupled to the locomotion speed of the mouse without any visual feedback coupling (**Methods**). The idea was to create an artificial closed loop coupling in which layer 2/3 now computes a difference between a motor-related prediction and a teaching signal it receives from the local layer 5 population. To test whether this was the case, we briefly broke the coupling between the optogenetic activation of layer 5 IT neurons and locomotion speed (**Figure 7A**) at random times for 1 s (akin to how we trigger visuomotor mismatch by halting visual flow) and measured neuronal responses to this perturbation. This optomotor mismatch resulted in strong responses in layer 2/3 that could not be explained by the offset of optogenetic stimulation alone (**Figure 7B**). The optomotor mismatch responses looked surprisingly similar to visuomotor mismatch responses on a population level (**Figure 7C**). Strikingly, by plotting the responses of all layer 2/3 neurons to optomotor mismatch against their responses to visuomotor mismatch, we found that there is a positive correlation between these two responses across the population, indicating that the same neurons are activated by both visuomotor and optomotor mismatch stimuli (**Figure 7D**). Thus, these data are consistent with the idea that the visuomotor mismatch responses we observe in layer 2/3 neurons are computed with layer 5 serving as a teaching signal, and that superficial cortical layers function to predict activity in deep cortical layers.

**Figure 7.**
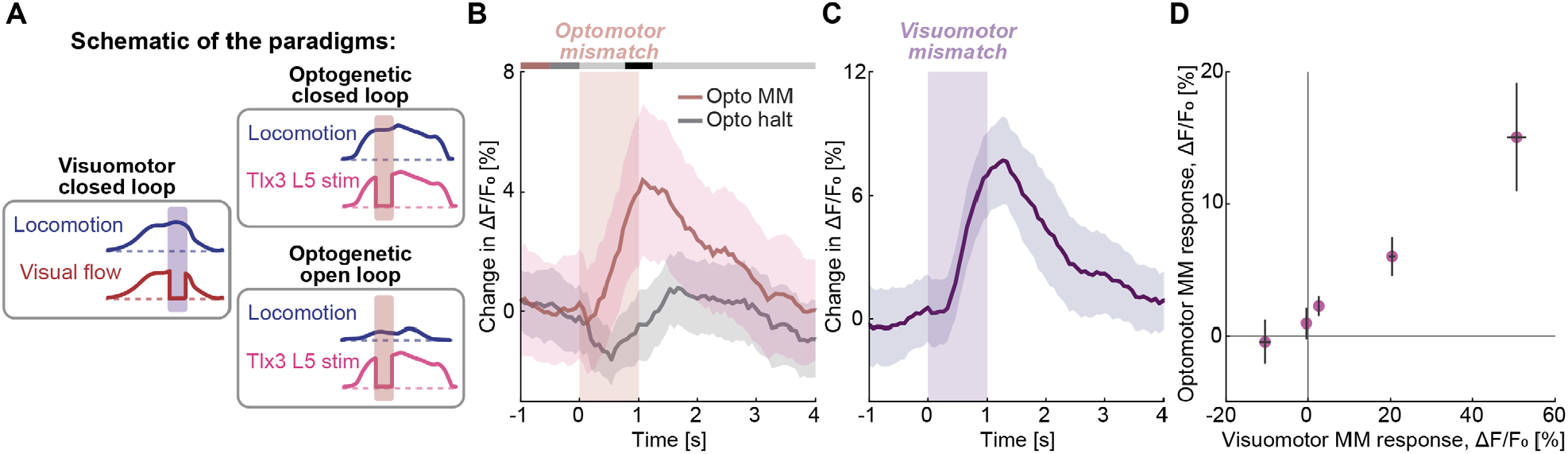
Layer 2/3 responds to manipulations of layer 5 activity as if layer 5 were a teaching signal for layer 2/3. **(A)** Mice first experienced a visuomotor closed loop condition, that was then followed by an optogenetic closed loop. In optogenetic closed loop, mice were exposed to constant visual illumination on the screen while the strength of optogenetic stimulation of Tlx3 layer 5 neurons was coupled to locomotion speed of the mouse instead. Following this, mice were exposed to an optogenetic open loop condition, in which the pattern of optogenetic stimulation the mouse had self-generated in the preceding optogenetic closed loop session was replayed. **(B)** Mean layer 2/3 population response to optomotor mismatch and optogenetic open loop stimulation halt. Pink shading indicates the duration of the stimulus. **(C)** Mean layer 2/3 population response to visuomotor mismatch. Purple shading indicates the duration of the visuomotor mismatch stimulus. **(D)** We binned layer 2/3 neurons by their visuomotor mismatch responses into 5 bins and plotted the mean optomotor mismatch responses of the neurons in these bins. Error bars indicate SEM.

Finally, if the input from deep cortical layers is the teaching signal for the prediction error computation in layer 2/3, what is the role of the visual input to layer 2/3 that arrives via layer 4? To begin addressing this question, we measured the functional influence from Scnn1a layer 4 neurons to layer 2/3 neurons (**Figure 8A**). Here, we found no evidence that the functional influence depended on whether a given layer 2/3 neuron was a positive or a negative prediction error neuron. (**Figures 8B-8F and S9**). Compared to functional influence of layer 5 IT, functional influence of layer 4 on layer 2/3 also did not depend on locomotion state of the mouse (**Figures S8G-S8I**). Thus, Scnn1a layer 4 neurons do not appear to function as a teaching signal for layer 2/3 prediction error neurons. Note, the Scnn1a-Cre line labels only one of the layer 4 cell types (Tasic et al., 2016), and there is direct thalamic input to layer 2/3 (Douglas et al., 1989; Morgenstern et al., 2016). While there may be other layer 4 cell types that function as teaching signal for layer 2/3, our findings indicate that the distinct modulation provided by layers 5 and 6 is not a universal property of all cortical inputs, but is likely less pronounced in circuits classically categorized as bottom-up.

**Figure 8.**
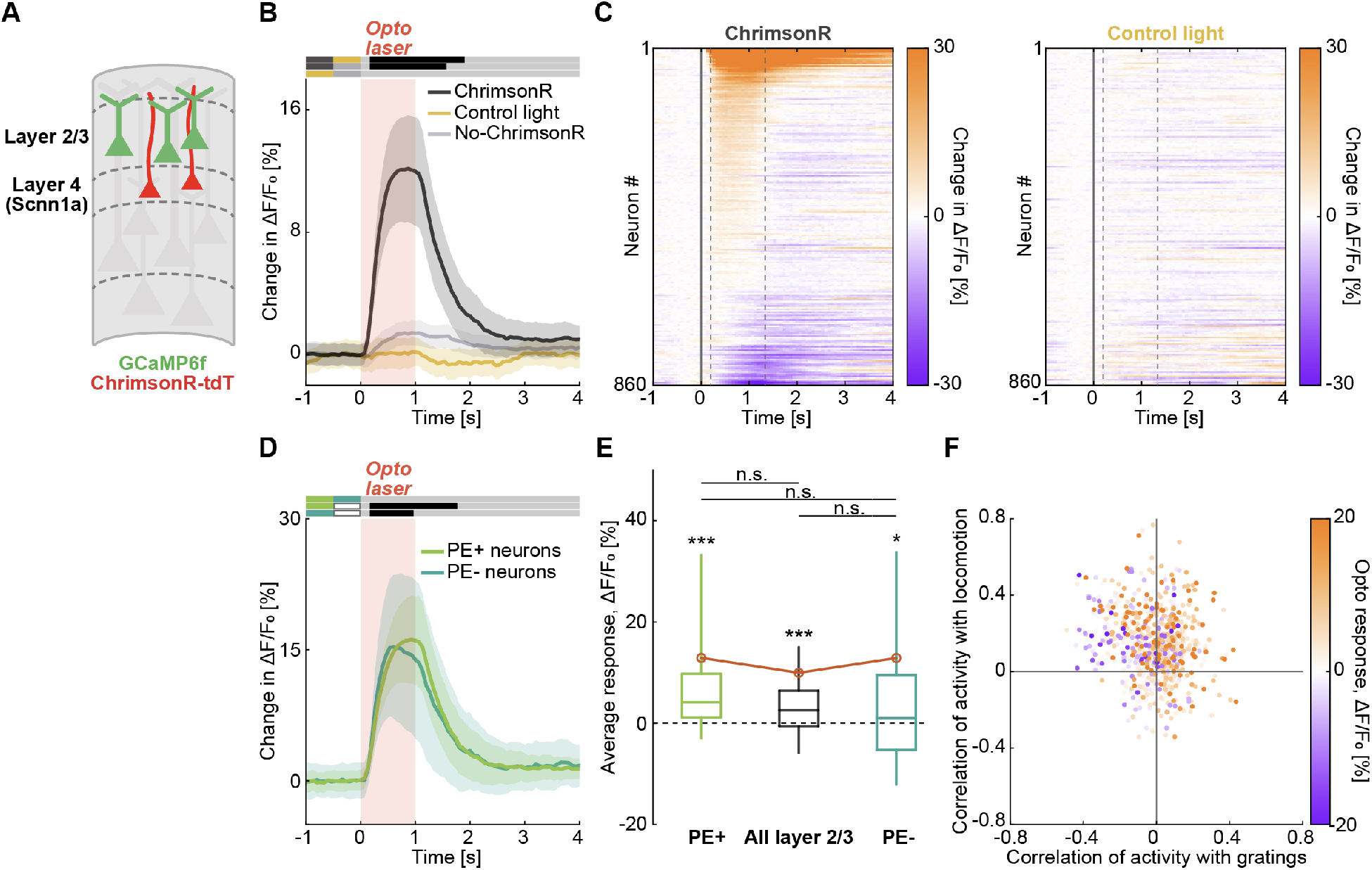
Functional influence of Scnn1a layer 4 neurons on positive and negative prediction error neurons of layer 2/3 is unspecific. **(A)** We optogenetically activated Scnn1a layer 4 neurons while recording calcium activity in layer 2/3 neurons. **(B)** Mean layer 2/3 population responses to optogenetic stimulation of Scnn1a layer 4 neurons (dark gray), control light stimulation in the same mice (yellow), and stimulation in no-ChrimsonR control mice (light gray). Red shading marks the stimulation interval. **(C)** Heatmap of mean layer 2/3 neuronal responses to optogenetic stimulation of Scnn1a layer 4 neurons (left) and control light stimulation (right). Neurons are sorted by the amplitude of their response to the optogenetic stimulation. **(D)** Mean responses of positive (light green) and negative (dark green) prediction error neurons in layer 2/3 to optogenetic stimulation of Scnn1a layer 4 neurons. Red shading marks the optogenetic stimulation interval. **(E)** Box plots showing average responses of positive prediction error neurons (light green), negative prediction error neurons (dark green), and all layer 2/3 neurons (black) to optogenetic stimulation of Scnn1a layer 4 neurons. Boxes indicate the interquartile range, the central line marks the median, and whiskers the 10th and 90th percentiles. Orange circles mark the mean response. **(F)** Scatter plot of the correlation between neuronal activity and optical flow speed against the correlation between neuronal activity and locomotion speed during grating sessions for all layer 2/3 neurons. Dot color indicates the amplitude of the response to optogenetic stimulation of Scnn1a layer 4 neurons.

## DISCUSSION

Here, we experimentally tested predictions of two circuit models of predictive processing (**Figure 1**) by mapping the functional influence between neurons of deep and superficial cortical layers. We found that functional influence from layers 5 and 6 on layer 2/3 (**Figures 3-5**) was consistent with the hierarchical predictive processing model but not with the non-hierarchical variant. Quite peculiarly, we also found that the functional influence in the reverse direction from layer 2/3 on layer 5 (**Figure 6**) was consistent with the non-hierarchical variant but not with the hierarchical one. Thus, under the assumption that prediction error neurons are located in superficial layers of cortex and internal representation neurons in deep layers, neither variant of the predictive processing model can account for the functional interactions between deep and superficial layers of cortex (**Figure S10A**).

### Limitations of the study

There are at least three caveats to our interpretation of the results:

1. The primary caveat is the partial screening coverage with respect to cell types. The mouse Cre lines and artificial regulatory elements we used to target excitatory neurons in layers 2/3, 4, 5, and 6 likely do not cover the space of all possible neuron types in these layers. In all cases, it is conceivable that there is a cell type we did not target that would have a functional influence profile opposite to what we have measured. In the case of the influence of layer 5 on layer 2/3 neurons, our experiments using layer 5 IT and layer 5 ET lines likely cover the majority of the dominant influence (Kim et al., 2015; Matho et al., 2021). In case of assessing the layer 4 influence on layer 2/3, the Scnn1a-Cre mouse only labels one of the subtypes of layer 4 excitatory neurons (Tasic et al., 2016). Here it is certainly possible that future experiments will find other layer 4 cell types with different profiles of functional influence on layer 2/3. However, our primary conclusions that layer 5 acts like a teaching input to layer 2/3 would not be changed by this. Finally, for the measurements of influence of layer 2/3 on layer 5, we have restricted our experiments to the influence on layer 5 IT neurons. This choice was motivated by the argument that layer 5 IT neurons are likely the primary target of layer 2/3 neurons in layer 5 (Anderson et al., 2010). Finally, it is likely that the two approaches used for targeting Tlx3 layer 5 neurons in this study (viral labelling versus a cross with a reporter line) may capture only partially overlapping subpopulations.
2. Another major caveat comes from the fact that we bulk stimulate entire populations of neurons. It is possible that each neuron exhibits a center-surround type interaction with its target population, such that a subset of neurons is excited while a larger set of off-target neurons is inhibited. If so, it is possible that in bulk stimulation experiments we would primarily observe the more dominant off-target interactions. If this were the case, all our conclusions on functional influence would be reversed. Assuming this applies to both layer 2/3 influence on layer 5 and vice versa, this would still mean that predictive processing cannot explain our results. It would however mean that layer 5 is not a teaching signal for layer 2/3, but a prediction. However, if our responses were indeed dominated by a surround effect, we would expect to find biphasic responses. Resolving this caveat will require an EM reconstruction approach (Schneider-Mizell et al., 2025).
3. Lastly, we cannot assess what fraction of the stimulation response can be explained by direct versus indirect interactions. For example, in the case of layer 5 stimulation, the influence on layer 2/3 is likely mediated by a combination of direct projections from layer 5 to layer 2/3, as well as via long-range cortical interactions, and subcortical loops. While, based on the relative strength, the reliability of responses, and the projection patterns between layer 5 and layer 2/3 (Hage et al., 2022), we speculate that the direct interaction is dominant, this is not of relevance to our computational interpretations.

### Joint embedding predictive architectures: a new starting point for understanding cortex?

There are two algorithms of cortical function we could identify that are consistent with our data. The first is a reciprocal variant of predictive processing, whereas the second is a joint embedding predictive architecture (JEPA). To illustrate the first, there are at least two ways in which predictive processing circuits can be arranged to exchange signals symmetrically. One is the non-hierarchical predictive processing variant we have tested here that is inconsistent with data. In this model both predictions and prediction errors are exchanged bidirectionally. For example, in the communication between visual and auditory cortices, auditory cortex sends a prediction of the visual teaching input to visual cortex given the current auditory representation and vice versa (Garner and Keller, 2022; Keller and Mrsic-Flogel, 2018). Similar arguments have been made for the interactions between sensory and motor areas (Jordan and Rumelhart, 1992; Keller and Sterzer, 2024). As we have established, this arrangement is inconsistent with our data. However, a computationally equivalent implementation would be one in which different cortical areas were to exchange predictions and teaching signals instead of predictions and prediction errors. This requires that each cortical area has two sets of prediction error neurons and two sets of corresponding internal representation neurons. If we add the condition that communication must be symmetrical, this would require two parallel prediction error sub-circuits in each node of the network, with two types of internal representation neurons. For these two sub-circuits the roles of teaching signal and prediction are reversed (**Figures S10B-S10C**), and so is the connectivity between layer 2/3 and layer 5. Thus, in such a model, we would expect to find both types of opposing influence in both directions in the interactions between layer 2/3 and layer 5. Thus, if we additionally assume that in our experiments, we have only identified parts of each sub-circuit (see caveats), our data would be consistent with this variant of non-hierarchical predictive processing. We speculate that such a reciprocal variant of predictive processing should be implementable given that a restricted hierarchical variant of this idea in the form of bidirectional predictive coding with shared representation neurons has been implemented (Oliviers et al., 2025).

However, we think the reciprocal variant of predictive processing is unlikely the correct model given recent evidence regarding the dominant dimension of coupling in cortex. All variants of predictive processing we discuss here, as well as all other models based on the idea of cortical columns, postulate a dominant role of vertical coupling in cortex: They predict that the activity in layer 5 of a cortical area is more strongly driven by the activity of layer 2/3 in that same area than by layer 5 activity in surrounding cortical areas. In predictive processing this comes from the assumption of a laminar separation between prediction error neurons and internal representation neurons. Such dominant vertical coupling is hard to reconcile with the fact that psychoactive drugs can selectively alter large scale activity patterns in layer 5 neurons of dorsal cortex without having corresponding effects in layer 2/3 (Bharioke et al., 2022; Heindorf and Keller, 2024; Vesuna et al., 2020). Thus, we conclude that no variant of predictive processing is consistent with both our data and a stronger horizontal than vertical coupling in cortex.

Based on these considerations, we here formulate a new testable hypothesis for the computational algorithm of cortex inspired by the idea of a JEPA (LeCun, 2022). Our proposal centers on the finding that layer 5 functionally interacts with layer 2/3 like a teaching input. Layer 5 input drives layer 2/3 prediction error neurons with a sign consistent with that expected of a teaching signal (**Figure 1**), and mismatches in the artificial coupling of layer 5 optogenetic activation to locomotion speed are sufficient to drive prediction error responses in layer 2/3 commensurate with those observed in visuomotor coupling (**Figure 7**). Thus, we propose that layer 2/3 functions to predict layer 5 activity, not sensory input per se, hence making predictions in the internal representation space, not input space. This highlights the key computational difference between JEPA and predictive processing. Predictive processing proposes formation of a generative internal model that constructs predictions in the input space. Consequently, prediction errors are also computed in the input space. In contrast, in a JEPA, predictions are formed in the latent space, and prediction error is computed and minimized in the latent space. According to predictive processing, we would expect the relevant teaching signal for layer 2/3 comparator to be bottom-up input into V1. However, our data instead argues that prediction errors are computed with respect to layer 5 activity as a teaching signal, which is more consistent with a JEPA. Combining this with the fact that deep and superficial layers of cortex each receive thalamic input (Constantinople and Bruno, 2013; Douglas et al., 1989), we speculate that cortex might function like a JEPA (Assran et al., 2023). A JEPA is a deep learning architecture that is designed to learn abstract representations of the input data in a self-supervised manner. It typically consists of two networks where a predictor uses representations of one network to predict the representation of the other (**Figure 9A**). The loss function in this architecture is based on the comparison between the predicted and the actual target representation. Classic instantiations of this principle in machine learning often use two identical or similar copies of encoder networks that employ some form of weight sharing. In the cortical network, however, it is difficult to conceptualize how such weight-sharing or copying would be implemented. In our analogy to cortex, we speculate that there are two separate networks that symmetrically serve as both encoder and target networks. Specifically, we propose that one of the two networks is housed in the superficial layers whereas the other resides in the deep layers (**Figure 9B**). The respective predictor networks connecting the two are implemented in columnar connectivity. Superficial and deep cortical layers embed a combination of thalamic and cortical input separately, and layer 2/3 prediction error neurons compute an error between the two embeddings. Thus, a JEPA-like architecture would also predict that the dominant intracortical communication driving activity within deep and superficial layers is horizontal (within layers) and not vertical (across layers), and would thus also simplify the explanation of how deep and superficial layers can be decoupled by psychoactive substances (Keller and Sterzer, 2024).

**Figure 9.**
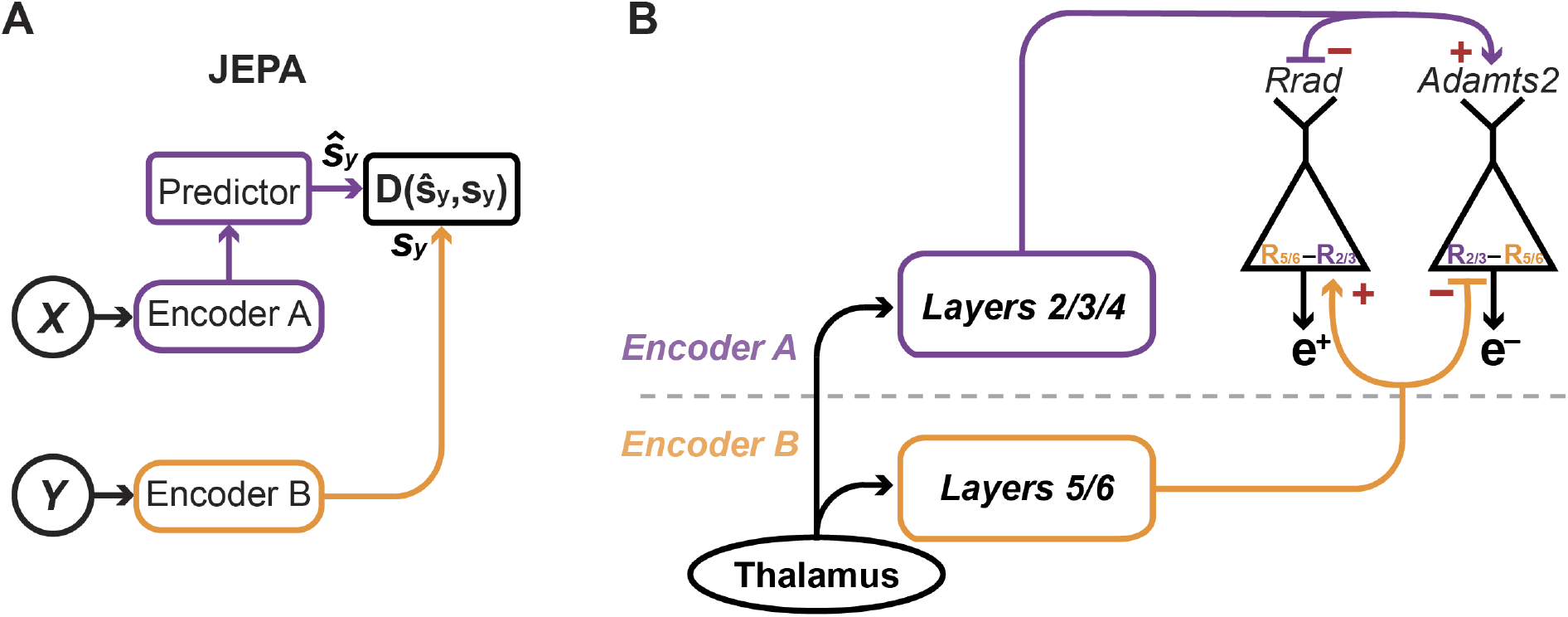
How a JEPA architecture could be mapped onto the cortical circuit. **(A)** Schematic of a JEPA architecture for self-supervised learning (adapted from Assran et al., 2023). The output of one encoder A is trained to predict the output of the other encoder B. The prediction is compared to the target output in a comparator network D. **(B)** A proposal for a JEPA implementation in cortical circuits. Superficial and deep cortical layers function as the two separate encoder networks postulated by JEPA. Prediction error neurons in layer 2/3 compare the local layer 5 activity to a prediction of that activity.

### Beyond JEPA

The core appeal of JEPA lies in its potential to perform a useful computation that can be mapped onto the local cortical microcircuit – a very promising, recently developed JEPA variant with an explicit analogy to cortex is recurrent predictive learning (RPL) (Mohammadi et al., 2025). It proposes two sequential networks, an encoder network whose output is fed to an integrator network. The output of the integrator network is then used to predict the output of the encoder, and the resulting prediction error drives learning. The encoder network would map onto superficial layers of cortex and the integrator network onto deep layers. Like a JEPA, RPL would also predict the existence of a prediction error computation that uses the encoder network (deep cortical layers) as a teaching signal.

However, a significant challenge remains: we have not found a way to map the algorithm onto the large-scale cortical architecture. The cortical network, as a whole, is fundamentally non-hierarchical. Most interactions in cortex are cross-modal, associative, or sensorimotor, and not hierarchical. This lack of a clear hierarchical arrangement is also apparent when the principles used to motivate the idea of a cortical hierarchy, based on anatomical data from the visual system (Felleman and Van Essen, 1991), are applied to data collected from across the whole cortex (Markov et al., 2013): Primary somatosensory cortex appears at the top of the “hierarchy”, primary visual cortex at the bottom. Likely, the only place where the hierarchical approximation is useful, is when analyzing an isolated, unimodal sensory processing stream. Thus, any serious candidate for an algorithm of cortical function must be implementable on a non-hierarchical network of nodes with symmetric interactions and arbitrary graph topology. JEPA – in its current variants – has been limited to hierarchical networks. We speculate that an interesting way forward would be to develop ways to implement a JEPA-like algorithm on a non-hierarchical network topology, as can be done for predictive processing (Salvatori et al., 2022). Inspired by JEPA and predictive processing, here we outline a set of ideas that that we think would constitute a foundation for a new model of cortical function. This proposal is informed by data, and our emphasis is on the elements that can be experimentally tested.

1. We propose that, similar to a JEPA, cortex is computationally split into two parallel networks embedded in deep, and superficial cortical layers respectively. Such a split is consistent with the dual-counterstream anatomy of cortex (Markov et al., 2013) and dense within layer connectivity (MICrONS Consortium, 2025). In this configuration, representations of superficial network are used to continuously predict activity in the deep cortical network to guide plasticity in deep layers.
2. Superficial cortical layers implement a predictive processing like algorithm with explicit prediction error neurons. However, instead of computing a difference between an external teaching input and a prediction, these error neurons compute a difference between the local representation in layer 5 and a prediction signal. Most prior evidence we have for predictive processing in cortex comes from evidence of prediction error responses in layer 2/3 and is consistent with this (Attinger et al., 2017; Fiser et al., 2016; Garner and Keller, 2022; Jordan and Keller, 2023, 2020; Keller et al., 2012; Leinweber et al., 2017; O’Toole et al., 2023; Solyga and Keller, 2025; Widmer et al., 2022; Zmarz and Keller, 2016). Based on this, we propose that two genetically identified subtypes of layer 2/3 neurons – Rrad and Adamts2 – function as positive and negative prediction error neurons respectively (O’Toole et al., 2023), while a third type of layer 2/3 neuron – Agmat – or a subset of layer 4 could function as internal representation neurons within superficial layers.
3. While the computational architecture of the deep cortical network is experimentally only poorly constrained, we speculate it could also be based on predictive processing. There is an intriguing mirroring of the layer 2/3 response profile in layer 5. Negative prediction error neurons are primarily found in superficial parts of layer 2/3 (O’Toole et al., 2023), likely what is layer 2 in higher mammals, while positive prediction error responses are enriched in deeper parts of layer 2/3. This is similar in layer 5 with strong visuomotor mismatch responses in the more superficial layer 5 IT population (Heindorf and Keller, 2024). It is conceivable that deep and superficial cortical layers might form a predictive processing like architecture, and that superficial layers implement predictive processing on latent representations of the deep layers. In this case superficial layers would constitute a predictive processing module that would be hierarchically above the deep layers, and would allow learning of multiple relational maps, e.g. learning of multiple task contingencies or generalizing across related tasks.
4. The source of external input (**Figure 9**) to both networks is thalamic projections. Both deep and superficial layers of cortex receive thalamic input (Constantinople and Bruno, 2013; Douglas et al., 1989). Theoretical work inspired by this arrangement also showed that such a network trained in a self-supervised manner would exhibit functional features consistent with those of cortical responses (Nejad et al., 2025).

The strength of this proposal lies in its integration of three key ideas: JEPA for self-supervised learning (Assran et al., 2023), predictive processing to solve the credit assignment problem (Whittington and Bogacz, 2017) and to enable learning on arbitrary graph topologies (Salvatori et al., 2022), and internal models for motor control (Jordan and Rumelhart, 1992; Kawato, 1999).

### Testable predictions

A useful model of a neuronal circuit is likely characterized by three principles:

1. The model is based on physiology and anatomy data.
2. It makes a concrete circuit proposal based on genetically identified cell types.
3. It can be implemented as an algorithm in silico to perform a useful computation.

These principles should enforce sufficient rigor to ensure that models are both informative of how the neuronal circuit might work and remain falsifiable. One of the few theories of cortical function that exhibited most of these characteristics is predictive processing. It was based on previously unexplained observations (Rao and Ballard, 1999), was consistent with the known cortical anatomy of the time (Bastos et al., 2012), there are concrete circuit proposals based on genetically identified cell types (Keller and Mrsic-Flogel, 2018), and it has some limited computational use when implemented (Lotter et al., 2016; Straka et al., 2020). However, given our results here, we conclude that predictive processing as formulated in its most prevalent variety (Keller and Mrsic-Flogel, 2018; Rao and Ballard, 1999) is inadequate to explain cortical computations.

The idea of a JEPA again appears like an opportune starting point to develop the next iteration of a hypothesis for the algorithm of cortex. It shares many of the circuit elements with predictive processing, which have been partially already confirmed to exist in layer 2/3, has computational usefulness, and would be consistent with our results here. There are several experimentally testable predictions we can make based on the idea that cortex implements a JEPA-like computation as detailed in the previous section:

1. Assuming superficial cortical layers implement one of two JEPA networks that only communicate with the network in deep cortical layers, but do not project directly out of cortex, then silencing layer 2/3 should interfere with learning in layer 5 but leave activity in layer 5, and behavior more generally, otherwise largely unaffected. This is in stark contrast to predictions the canonical microcircuit model would make.
2. A JEPA would also predict a difference in learning rates between the encoder networks and the predictor network (Assran et al., 2023; Grill et al., 2020; Mohammadi et al., 2025). Based on this, we should find higher levels of plasticity in the layer 2/3 prediction error network, than in layer 4 or the corresponding representation network in deep cortical layers.
3. There should be a neuron type in layer 2/3 or layer 4 that acts as an internal representation neuron for the prediction error neurons in layer 2/3. This could either be Agmat positive layer 2/3 neurons (O’Toole et al., 2023), or a subset of layer 4 neurons. These neurons should exhibit opposing functional influence patterns on prediction error neurons as predicted by the predictive processing circuit model (Keller and Mrsic-Flogel, 2018).

We believe that testing predictions outlined above is a promising way forward to further constrain the search for a candidate algorithm of cortical function and formulate a more complete proposal.

## METHODS

### Animals

All animal procedures were approved by and carried out in accordance with guidelines of the Veterinary Department of the Canton Basel-Stadt, Switzerland. The mice used in this study were: 23 Tlx3-Cre (Gerfen et al., 2013), 6 Fezf2-CreERT2 (Matho et al., 2021), 6 Ntsr1-Cre (Gong et al., 2007), 4 Scnn1a-Cre (Madisen et al., 2010) and 8 Tlx3-Cre x Ai148 (Daigle et al., 2018). Transgenic mice were bred in-house and heterozygous mice of either sex were used. 8 females of C57BL/6J (Charles River Laboratories) were used for control experiments. All mice were aged between 7 and 22 weeks. Mice were co-housed in groups of 2 to 5 in a reversed 12 hour light-dark cycle and had ad libitum access to water and regular mouse food. To induce expression of Cre in Fezf2-CreERT2 mice, their regular food was substituted for tamoxifen containing food (Envigo, 400 mg tamoxifen per kg of food) for the duration of 3 weeks. All imaging experiments took place in the dark phase of the light cycle.

### Surgery and viral vector injections

For all surgical procedures, mice were anesthetized using a mix of fentanyl (0.05 mg/kg), medetomidine (0.5 mg/kg) and midazolam (5 mg/kg). Analgesics were applied peri- and postoperatively. Lidocaine was injected locally on the scalp (10 mg/kg s.c.) prior to surgery. To provide postoperative analgesia we used either a combination of repeated injections of metacam (5 mg/kg, s.c.) and buprenorphine (0.1 mg/kg s.c.) or extended-release buprenorphine (Ethiqa XR, 3.25 mg/kg s.c.). Mice were returned to their home cage after the anesthesia was antagonized by a mix of flumazenil (0.5 mg/kg) and atipamezole (2.5 mg/kg) and were allowed to recover for at least one week prior to first head-fixation.

For two-photon imaging and concurrent optogenetic stimulation experiments, mice underwent a cranial window implantation surgery. First, a circular craniotomy 4 mm in diameter was made over right V1 centered on 2.5 mm lateral and 0.5 mm anterior of lambda, which was followed by a durectomy. Then an AAV vector mix was injected (approximately 300 nl per injection) across 4-6 locations spanning right V1. For mapping the functional influence from cell types of deep layers on layer 2/3, the AAV vector mix consisted of AAV2/1-EF1α-GCaMP6f (5×10^12^ - 4×10^14^ GC/ml) and AAV2/1-hSyn-DIO-ChrimsonR-tdTomato (10^13^ – 3×10^14^ GC/ml). For no-ChrimsonR control mice, the ChrimsonR vector was omitted. For mapping the functional influence from cell types of layer 2/3 onto layer 5, Tlx3-Cre x Ai148 mice were injected with either AAV2/1-AP.Baz1a.1-CreOFF-ChrimsonR-mRuby-ST (4.6×10^13^ GC/ml) or AAV2/1-AP.Adamts2.1-CreOFF-ChrimsonR-mRuby-ST (2.5×10^13^ GC/ml). For assessing the responses of Tlx3 neurons to optogenetic activation of the same population, mice were injected with AAV2/9-DIO-jGCaMP8s-P2A-ChrimsonR-ST (1.1×10^15^ GC/ml). A 4 mm diameter glass window was then placed over the craniotomy and sealed with cyanoacrylate. Lastly, a custom titanium headplate was attached to the skull using dental cement (Paladur, Heraeus).

### Virtual reality and visuomotor paradigms

During all recordings, mice were head-fixed in a virtual reality system as described previously (Leinweber et al., 2014). Briefly, mice were free to run on an air-supported polystyrene ball of 20 cm in diameter, and the virtual environment was projected onto a toroidal screen surrounding the mouse. From the point of view of the mouse, the screen covered a visual field of approximately 240 degrees horizontally and 100 degrees vertically. The projector output was gated by a 24 kHz gating signal to only turn the illumination on at the turnaround points of the two-photon resonance scanner. This was done to minimize light leak from the virtual reality system. In the closed loop and open loop conditions, the virtual environment presented on the screen was a virtual tunnel with walls textured with vertical sinusoidal gratings. Movement in the virtual tunnel was restricted to one dimension (forward and backwards). In the closed loop condition, visual flow speed experienced by the mouse was coupled to its locomotion speed, with the exception of brief (1 s) mismatch events presented every 12 ± 4.5 s (mean ± SD). During these events visual flow speed was clamped to zero for 1 s. In the open loop condition, the visual flow presented was a playback of the visual flow the mouse had self-generated in the preceding closed loop session. For the grating sessions, we presented the mice with full-field sinusoidal drifting gratings at 0 degrees, 45 degrees, 90 degrees and 135 degrees orientations, moving in either direction. Each stimulus in a sequence was selected randomly and presented for a duration of 6 ± 1.35 s (mean ± SD) with a randomized inter-stimulus interval of 4.5 ± 1 s (mean ± SD) during which a gray screen was displayed. For closed loop and open loop optogenetic activation of layer 5 experiments (**Figure 7**), mice were presented with a static gray screen. Prior to the imaging experiments, mice were habituated to head-fixation and locomotion on the ball during daily 1-hour training sessions for at least 3 days, until they displayed regular locomotion.

### Two-photon calcium imaging

Two-photon calcium imaging was performed using a modified Thorlabs Bergamo II two-photon microscope. Excitation illumination was provided by a titanium-sapphire laser (Mai Tai, Spectra-Physics) tuned to 930 nm. The emission light was band-pass filtered using a 525/50 filter (Semrock) and detected by a photomultiplier tube (H7422, Hamamatsu). The detected signal was amplified (DHPCA-100, Femto), digitized (NI5772, National Instruments) and band-pass filtered at 80 MHz using a digital Fourier-transform filter implemented on an FPGA (NI5772, National Instruments) using custom-written software. A 12 kHz scanning system (Cambridge Technology) was used to acquire images at 750 x 400 pixels resolution at 60 Hz frame rate. The field of view under the microscope was approximately 300 x 300 µm. A piezo electric linear actuator (P726 PIFOC, Physik Instrumente) was used to move the objective (Nikon 16x, 0.8 NA) to image 4 separate planes sequentially, which resulted in an effective frame rate of 15 Hz. Custom software was used to acquire imaging data.

### Concurrent two-photon imaging and optogenetic stimulation

Illumination source for the optogenetic stimulation of ChrimsonR-expressing neurons was a 637 nm laser (OBIS, Coherent). The laser beam was merged with the main imaging path using a dichroic mirror (ZT775sp-2p, Chroma) and sent through the two-photon microscope objective. Another dichroic mirror (F38-555SG, Semrock) was used to split the GFP emission from both illumination light sources. To reduce the light leak artifacts during optogenetic stimulation, the laser output was synchronized to the turnaround times of the resonant scanner. An essential addition to artifact free stimulation was the digital band-pass filter used to process the PMT signals (see above). To optogenetically activate cell types of layers 4, 5 and 6 (**Figures 3-5 and 8**), we used contentious laser pulses of 1 s duration, presented randomly in closed loop and open loop sessions every 12 ± 3.5 s. The laser power used for optogenetic stimulation in these experiments was fixed at 11 mW. For optogenetic activation of layer 2/3 cell types (**Figure 6**) stimulation occurred on average once every 11 s, and the stimulation power varied between 1 mW and 11 mW. The responses were then averaged over all stimulation powers.

### Closed loop and open loop optogenetic activation of layer 5

For the closed loop optogenetic activation experiment (**Figure 7**) the stimulation laser power was coupled to the locomotion speed of the mouse. The stimulation power scaled linearly with the speed of the mouse and ranged from the minimum of 0 mW (when mice were stationary), to a maximum of 6 mW (set to the maximum locomotion speed of the preceding sessions). The opto-motor mismatch events were triggered by clamping the stimulation power to 0 mW for 1 s at random times every 12 ± 3.5 s (mean ± SD). The open loop condition consisted of a replay of the stimulation power profile the mouse had self-generated in the preceding closed loop session including optomotor mismatch events.

### Extraction of neuronal activity

Functional two-photon calcium imaging data were processed as previously described (Keller et al., 2012). Briefly, raw images were full-frame registered to correct for lateral brain motion. Neurons were selected manually based on mean and maximum fluorescence images. Fluorescence traces of each neuron were calculated as the mean fluorescence over all pixels in the given region of interest and corrected for slow drift in fluorescence using an 8^th^-percentile filter and 1000 frame (67 s) window. For ΔF/F_0_ calculation, F_0_ was defined as the median fluorescence over the entire trace.

### Data analysis

To quantify the average response traces, we used a hierarchical bootstrap approach (Saravanan et al., 2020) to estimate the mean activity value at each time bin. Briefly, for each trace the data were first resampled with replacement at the level of recording sites. From the sampled set of recording sites, the data were then resampled with replacement at the level of neurons. For each of 10 000 such bootstrap samples we calculated the mean, to generate a distribution of bootstrap mean values. We used the mean of this distribution as our estimate of the true mean, and the standard deviation of this distribution as bootstrap standard error. All average response traces were then baseline (mean ΔF/F_0_ in 0.6 s window prior to the stimulus onset) subtracted.

For the analysis of visuomotor and optomotor mismatch responses, we included only mismatch events that occurred during locomotion periods – when the average locomotion velocity in the window -1 to +1 s around the mismatch onset was above 2.7 cm/s. Visuomotor mismatch events that overlapped with optogenetic stimulation in the interval of -1 to +1 s from mismatch onset were excluded from the analysis. For the analysis of grating onset responses, we included only grating onsets that occurred during stationary periods (when the average locomotion velocity in the -1 to +1 s window around grating onset was below 1.8 cm/s). The only exception was the analysis of responses to a specific grating orientation (**Figures S5A and S5B**), for which all trials were included due to a low number of repetitions per orientation. The same windows and thresholds were used to classify locomotion and stationary periods for the analysis of optogenetic stimulation responses (**Figure S8**). Locomotion onset was defined as a time point at which locomotion speed crossed a threshold of 1.8 cm/s, provided that the mean speed in the preceding 1 s was below the threshold and the mean speed in the following 1 s remained above this threshold. For analysis of locomotion onsets during open loop or grating sessions, locomotion onsets that occurred within a window -0.33 to + 2 s relative to a visual stimulus onset were excluded.

Neuronal responses to visuomotor mismatch, grating onset, optomotor mismatch and optomotor open loop stimulation offset were calculated as the mean activity of each neuron in a 0.33 to 1.33 s response window after stimulation onset minus the mean activity of each neuron in a 0.6 s baseline window preceding the stimulation onset. For quantification of neuronal responses to optogenetic activation, a response window of 0.2 to 1.33 s was used to account for faster response dynamics. To identify positive and negative prediction error neurons, neurons were classified based on their responses to grating onset and visuomotor mismatch, respectively. For each stimulus, the top 15% (**Figures 2-5, 8, S1, S3-S7, S9**), 10% or 20% (**Figures S3, S6, S7, S9**) of responsive neurons were selected ensuring no overlap. For all analyses in which we select and plot on the same variable (**Figures 2-6, 8, S5**), we selected neurons based on half the trials and used the other half of the trials for plotting. This was done to avoid one type of regression to the mean artifact that results from selecting the most responsive neurons to a given stimulus and then plotting the responses of these neurons based on the same data. To prevent graphical aliasing, all heatmaps were smoothed across 3 to 30 neurons, with the smoothing window scaled to the size of the plotted population to ensure correct display at minimum resolution of 210 pixels per heatmap. To determine the correlation between neuronal activity and optical flow speed or locomotion speed during the grating sessions, we calculated the correlation coefficient between the individual neuronal activity traces and binarized grating trace or activity traces and locomotion speed respectively (**Figures 2-5 and 8**). To quantify the neuronal responses to optomotor mismatch as a function of visuomotor mismatch response (**Figure 7**), we binned layer 2/3 neurons in 5 groups based on their visuomotor mismatch response. The bins comprised 0th-10th, 10th-25th, 25th-85th, 85th-95th and 95th-100th percentiles of responses distribution. The top two bins were chosen to reflect the top 15% of neurons used for analysis elsewhere, the rest were chosen pseudo randomly. Exact choice of bins did not influence our conclusions.

### Statistical tests

All statistical information for the tests performed in this manuscript is provided in **Table S1**. For statistical testing we used a hierarchical bootstrapping approach to account for the nested structure of the data. Each recording site contains a different number of neurons, so neurons (lower level of the hierarchical data structure) are grouped within the recording sites (higher level of the hierarchical data structure). Data are resampled hierarchically to generate bootstrap estimates (Saravanan et al., 2020). For the quantification of average response traces, the data were first resampled with replacement at the level of recording sites and then resampled with replacement at the level of neurons. For each of 10 000 such bootstrap samples we calculated the mean to generate a bootstrap distribution of mean values. The p-value was then calculated as a fraction of the bootstrap sample means which violated the tested hypothesis. To report the difference between two average calcium traces (**Figures 2-8, S1, S3-S9**), we performed hierarchical bootstrapping for every time bin of the calcium trace and marked bins where responses were significantly different (p < 0.05). We removed isolated significant bins, such that only intervals of at least two consecutive significant bins were marked.

## Data and software availability

All raw data and software to generate all figures in this manuscript will be deposited to zenodo.org upon submission as the Version of Record. Software to control the two-photon microscope is available at https://sourceforge.net/projects/iris-scanning/.

## ACKNOWLEDGEMENTS

We thank Ashena Gorgan Mohammadi, Manu Srinath Halvagal, and Friedemann Zenke for inspiration and discussion as well as comments on earlier versions of this manuscript. We thank Tingjia Lu for production of the AAV vectors, Jennifer Gröli for animal husbandry, and Z. Josh Huang for sharing Fezf2-CreERT2 mice. We have received generous donations of Dr. Pepper from Felix Widmer. We thank all the members of the Keller lab for discussion and support. This project has received funding from the Swiss National Science Foundation (GBK), the Novartis Research Foundation (GBK), and the European Research Council (ERC) under the European Union’s Horizon 2020 research and innovation programme (grant agreement No 865617) (GBK).

## AUTHOR CONTRIBUTIONS

AV designed and performed the experiments and analyzed the data. All authors wrote the manuscript.

## SUPPLEMENTARY FIGURES

**Figure S1.**
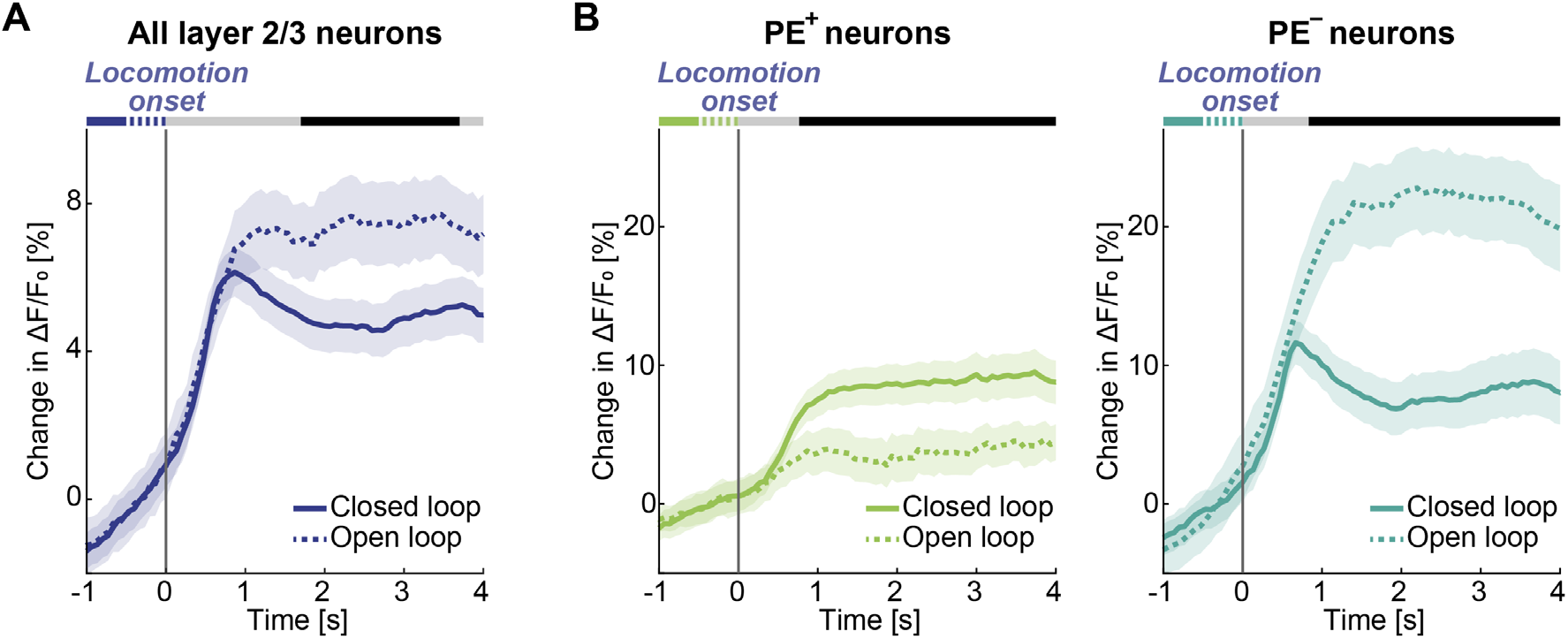
Locomotion onset responses of layer 2/3 neurons differ between closed loop and open loop conditions. **(A)** Mean layer 2/3 population response to locomotion onset in closed loop (solid line) and open loop (dashed line) conditions. Here and in the subsequent panels, response curves represent the hierarchical bootstrap estimates of the mean trace. Shading around the line is the bootstrap error, defined as one standard deviation of the bootstrap distribution at each time bin. Response curves are compared for each time bin: in the horizontal bars above the plot, black marks time bins where p<0.05 and gray mark time bins where p>0.05. **(B)** Mean response of positive (left, light green) and negative (right, dark green) prediction error neurons to locomotion onset in closed loop (solid lines) and open loop (dashed lines) conditions.

**Figure S2.**
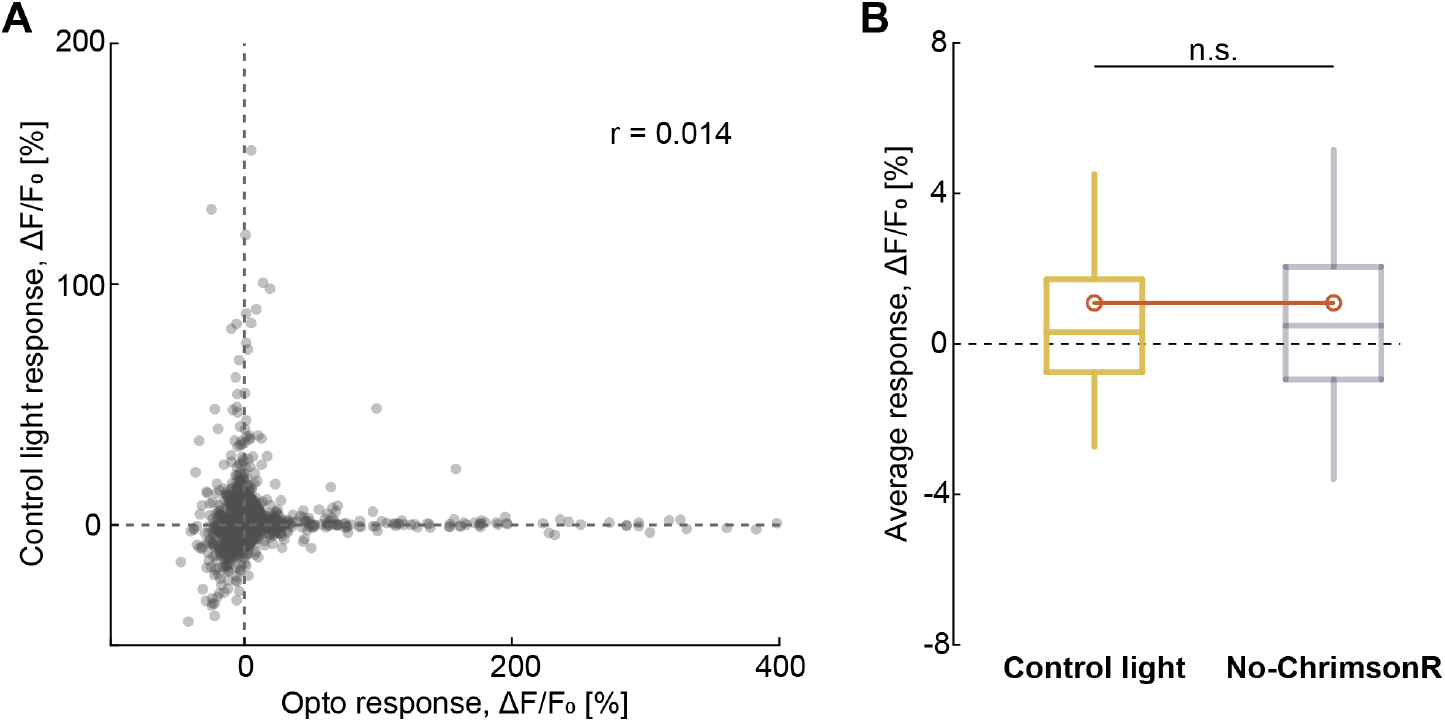
Responses of layer 2/3 neurons to control light stimulation do not correlate with optogenetic stimulation responses and resemble responses in no-ChrimsonR control mice. **(A)** Scatterplot of the responses of layer 2/3 neurons to optogenetic stimulation of Tlx3 layer 5 neurons against their responses to control light stimulation. Each dot represents a neuron. **(B)** Comparison of average responses of layer 2/3 neurons to control light stimulation (yellow) and stimulation light in no-ChrimsonR control mice (light gray). Boxes indicate the interquartile range, the central line marks the median, and whiskers the 10th and 90th percentiles. Orange circles mark the mean response.

**Figure S3.**
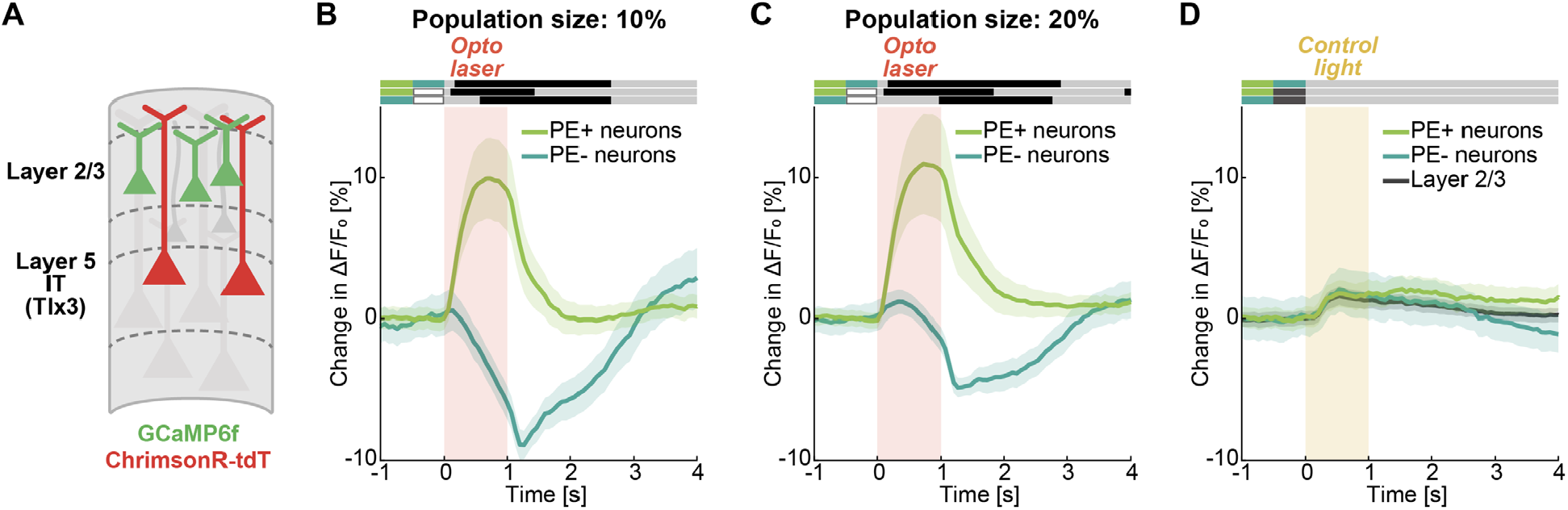
Functionally specific modulation by Tlx3 layer 5 neurons is robust to functional group size and is absent for control light stimulation. **(A)** We optogenetically activated Tlx3 layer 5 neurons while recording calcium activity in layer 2/3 neurons. **(B)** As in **Figure 3D**, but using the 10% (instead of 15%) most strongly responding neurons to grating onset and visuomotor mismatch. **(C)** As in **Figure 3D**, but using the 20% (instead of 15%) most strongly responding neurons to grating onset and visuomotor mismatch. **(D)** Mean responses of positive prediction error neurons (light green), negative prediction error neurons (dark green), and the layer 2/3 population (black) to control light stimulation. Yellow shading marks the control stimulation interval.

**Figure S4.**
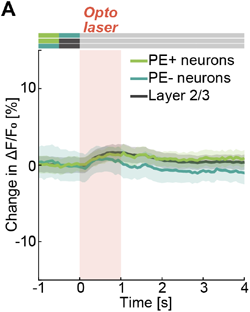
No-ChrimsonR control stimulation does not induce functionally specific modulation of layer 2/3 neurons. (**A**) Mean responses of positive prediction error neurons (light green), negative prediction error neurons (dark green), and the layer 2/3 population (black) to stimulation light in no-ChrimsonR control mice. Red shading marks the stimulation interval.

**Figure S5.**
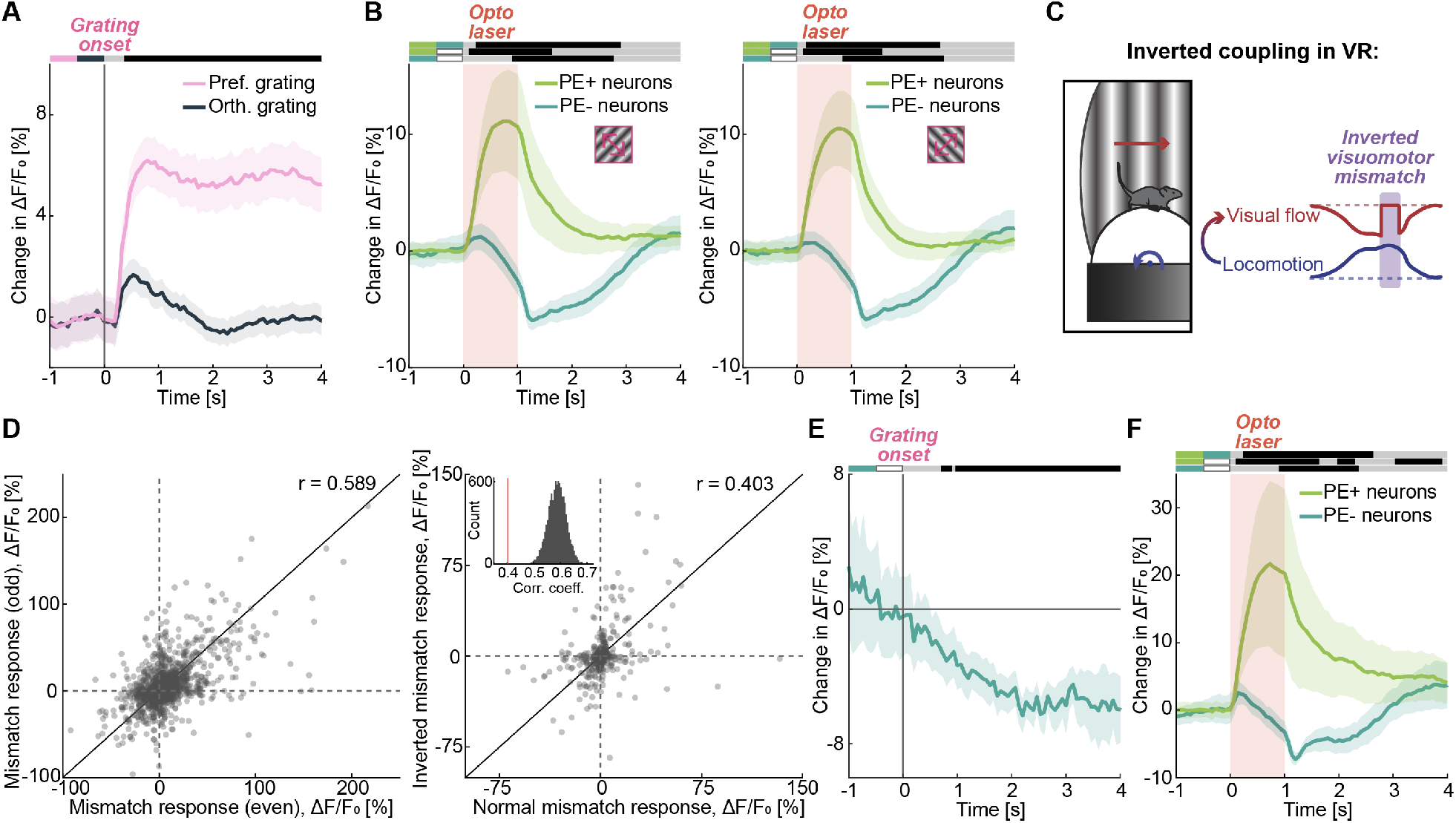
Stimulation of layer 5 likely separates positive and negative error neurons irrespective of feature-selectivity of those neurons. **(A)** Mean layer 2/3 population response to preferred grating (pink) and orthogonal grating (black) onsets. **(B)** Mean responses of positive prediction error neurons (light green) and negative prediction error neurons (dark green) to optogenetic stimulation of Tlx3 layer 5 neurons. Positive prediction error neurons were selected using a single grating orientation, either 135° (left) or 45° (right). Red shading marks the stimulation interval. **(C)** Schematic of the inverted visuomotor coupling in the virtual reality system. Locomotion of the mouse on the treadmill is coupled to forward moving visual flow. We define inverted visuomotor mismatch stimulus as a halt in the inverted visual flow during locomotion of the mouse. **(D)** Inverted visuomotor mismatch drives a population of neurons different from those activated by normal visuomotor mismatch. Left: quantification of measurement noise in normal visuomotor mismatch responses. Scatter plot of visuomotor mismatch response on even trials versus the same response on odd trials. Each dot is a neuron. The black line marks the diagonal. Right: Responses of layer 2/3 neurons to normal visuomotor mismatch versus their responses to inverted visuomotor mismatch. The correlation between the two is significantly lower than would be expected from measurement noise. Inset: Bootstrap distribution of odd versus even trial correlations of normal visuomotor mismatch responses. Vertical red line marks the observed correlation between normal and inverted visuomotor mismatch responses. **(E)** Mean response of inverted visuomotor mismatch neurons to grating onset. Like normal visuomotor mismatch neurons (**Figure 2F**), inverted visuomotor mismatch neurons decrease their activity on visual stimulation. **(F)** Mean responses of positive prediction error neurons (light green) and negative prediction error neurons (dark green) to optogenetic stimulation of Tlx3 layer 5 neurons. Negative prediction error neurons are selected using inverted visuomotor mismatch stimulus. Red shading marks the stimulation interval.

**Figure S6.**
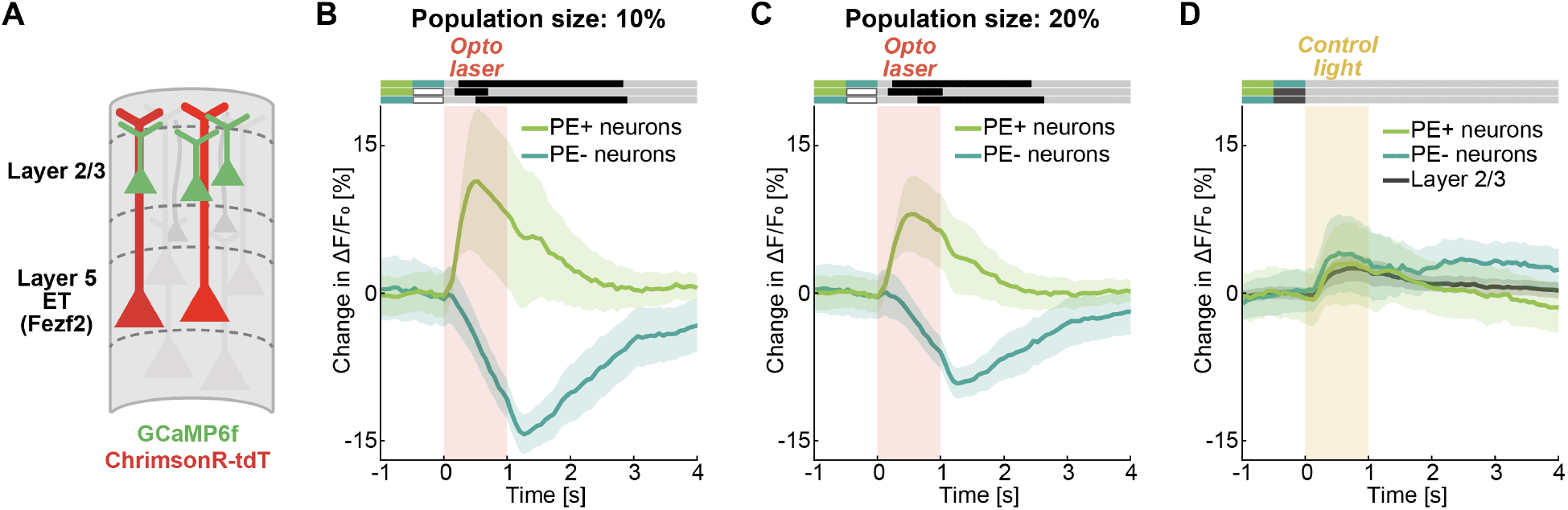
Functionally specific modulation by Fezf2 layer 5 neurons is robust to functional group size and is absent for control light stimulation. **(A)** We optogenetically activated Fezf2 layer 5 neurons while recording calcium activity in layer 2/3 neurons. **(B)** As in **Figure 4D**, but using the 10% (instead of 15%) most strongly responding neurons to grating onset and visuomotor mismatch. **(C)** As in **Figure 4D**, but using the 20% (instead of 15%) most strongly responding neurons to grating onset and visuomotor mismatch. **(D)** Mean responses of positive prediction error neurons (light green), negative prediction error neurons (dark green), and the layer 2/3 population (black) to control light stimulation. Yellow shading marks the control stimulation interval.

**Figure S7.**
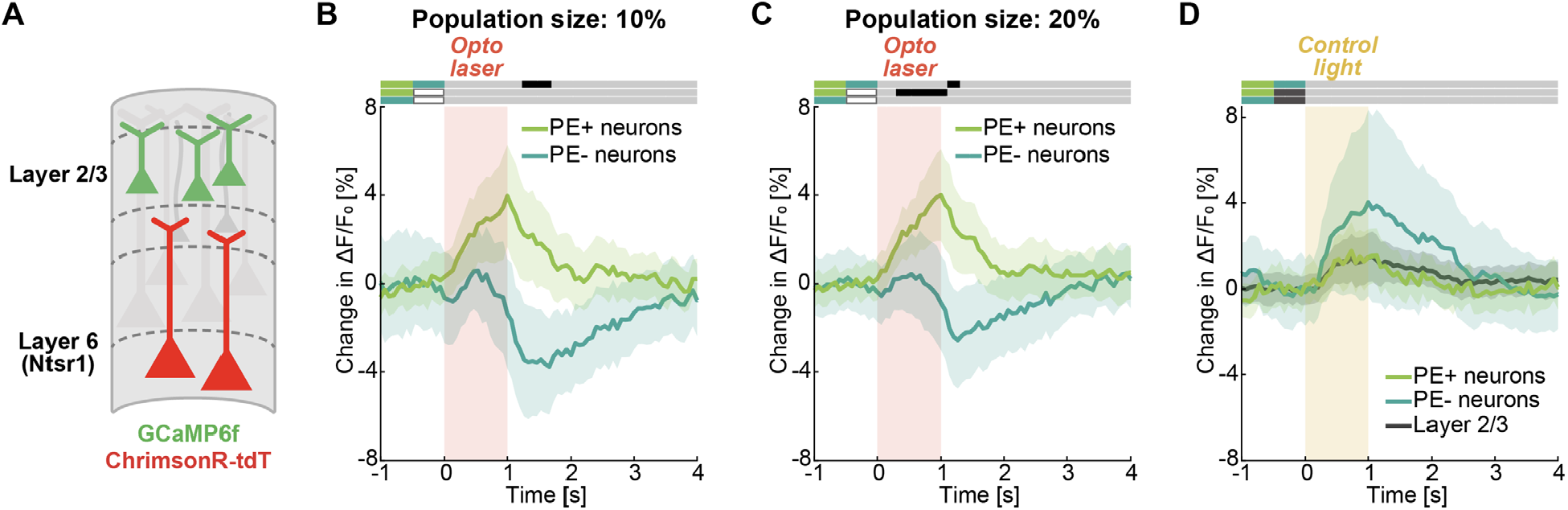
Functionally specific modulation by Ntsr1 layer 6 neurons is robust to functional group size and is absent for control light stimulation. **(A)** We optogenetically activated Ntsr1 layer 6 neurons while recording calcium activity in layer 2/3 neurons. **(B)** As in **Figure 5D**, but using the 10% (instead of 15%) most strongly responding neurons to grating onset and visuomotor mismatch. **(C)** As in **Figure 5D**, but using the 20% (instead of 15%) most strongly responding neurons to grating onset and visuomotor mismatch. **(D)** Mean responses of positive prediction error neurons (light green), negative prediction error neurons (dark green), and the layer 2/3 population (black) to control light stimulation. Yellow shading marks the control stimulation interval.

**Figure S8.**
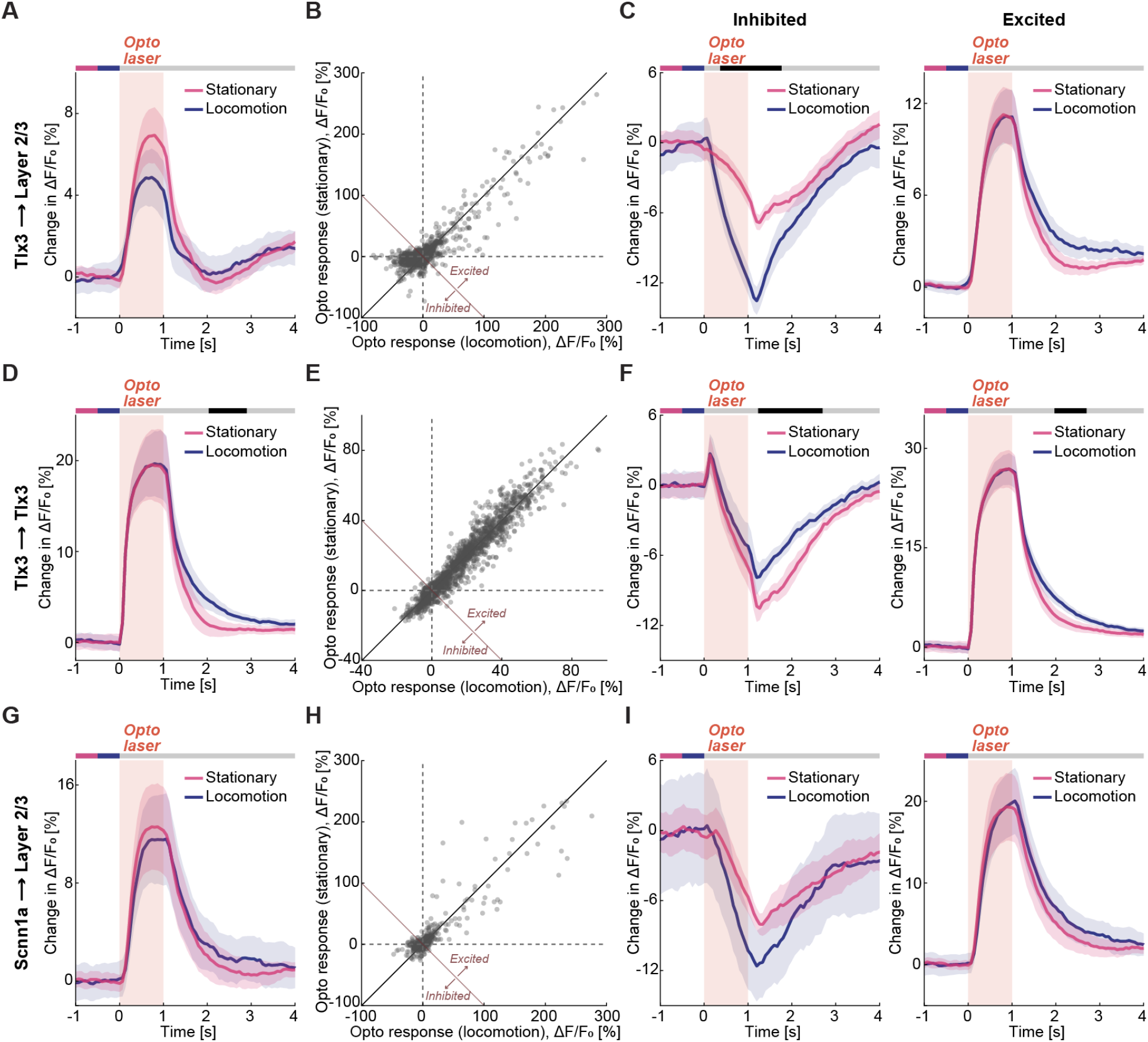
Locomotion unmasks inhibitory influence of layer 5 IT neurons, but not layer 4 neurons, on layer 2/3 neurons. **(A)** Mean layer 2/3 population response to optogenetic stimulation of Tlx3 layer 5 neurons split by whether the mouse is stationary (pink) or locomoting (blue). **(B)** Scatterplot of layer 2/3 neuronal responses to stimulation of Tlx3 layer 5 neurons while mice were locomoting versus their responses while mice were stationary. Each dot is a neuron. For the analysis shown in **C**, we define neurons below and to the left of the negative diagonal as inhibited, and those to the right and above as excited. **(C)** Left: Mean response of inhibited layer 2/3 neurons (as defined in **B**). Right: Mean response of excited layer 2/3 neurons. (**D-F**) As in **A-C**, but for responses of Tlx3 layer 5 neurons to optogenetic stimulation of the same population. (**G-I**) As in **A-C**, but for responses of layer 2/3 neurons to optogenetic stimulation of Scnn1a layer 4 neurons.

**Figure S9.**
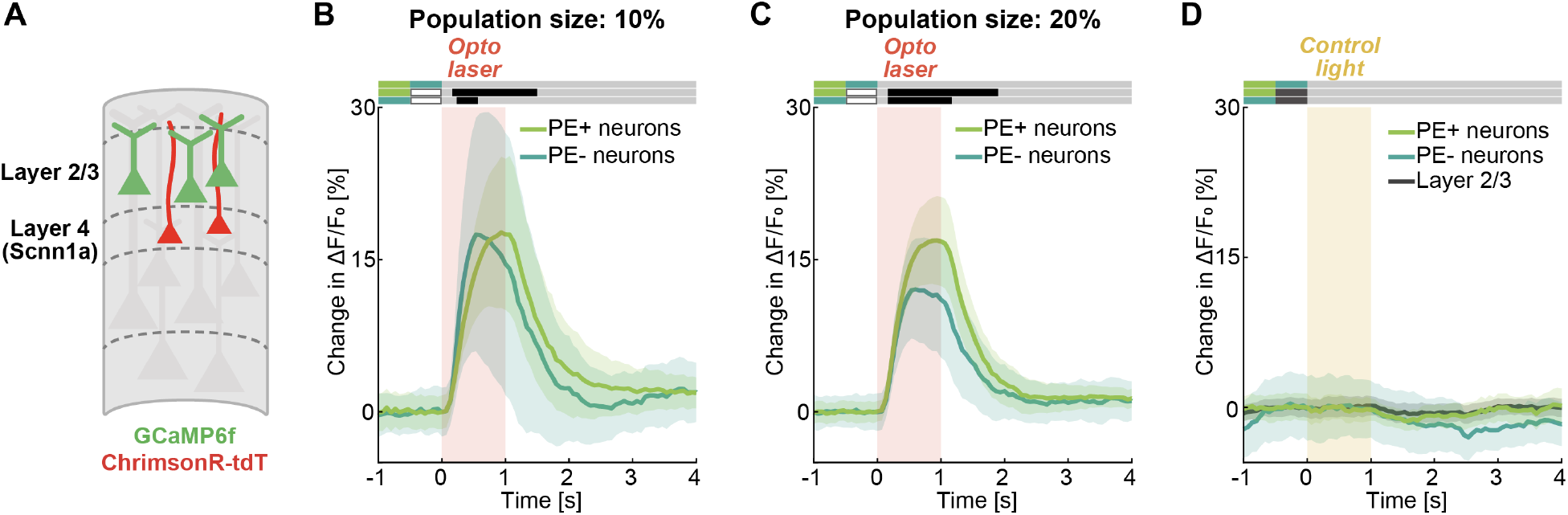
Neither optogenetic stimulation of Scnn1a layer 4 neurons nor control light stimulation results in functionally specific modulation in layer 2/3. **(A)** We optogenetically activated Scnn1a layer 4 neurons while recording calcium activity in layer 2/3 neurons. **(B)** As in **Figure 8D**, but using the 10% (instead of 15%) most strongly responding neurons to grating onset and visuomotor mismatch. **(C)** As in **Figure 8D**, but using the 20% (instead of 15%) most strongly responding neurons to grating onset and visuomotor mismatch. **(D)** Mean responses of positive prediction error neurons (light green), negative prediction error neurons (dark green), and the layer 2/3 population (black) to control light stimulation. Yellow shading marks the control stimulation interval.

**Figure S10.**
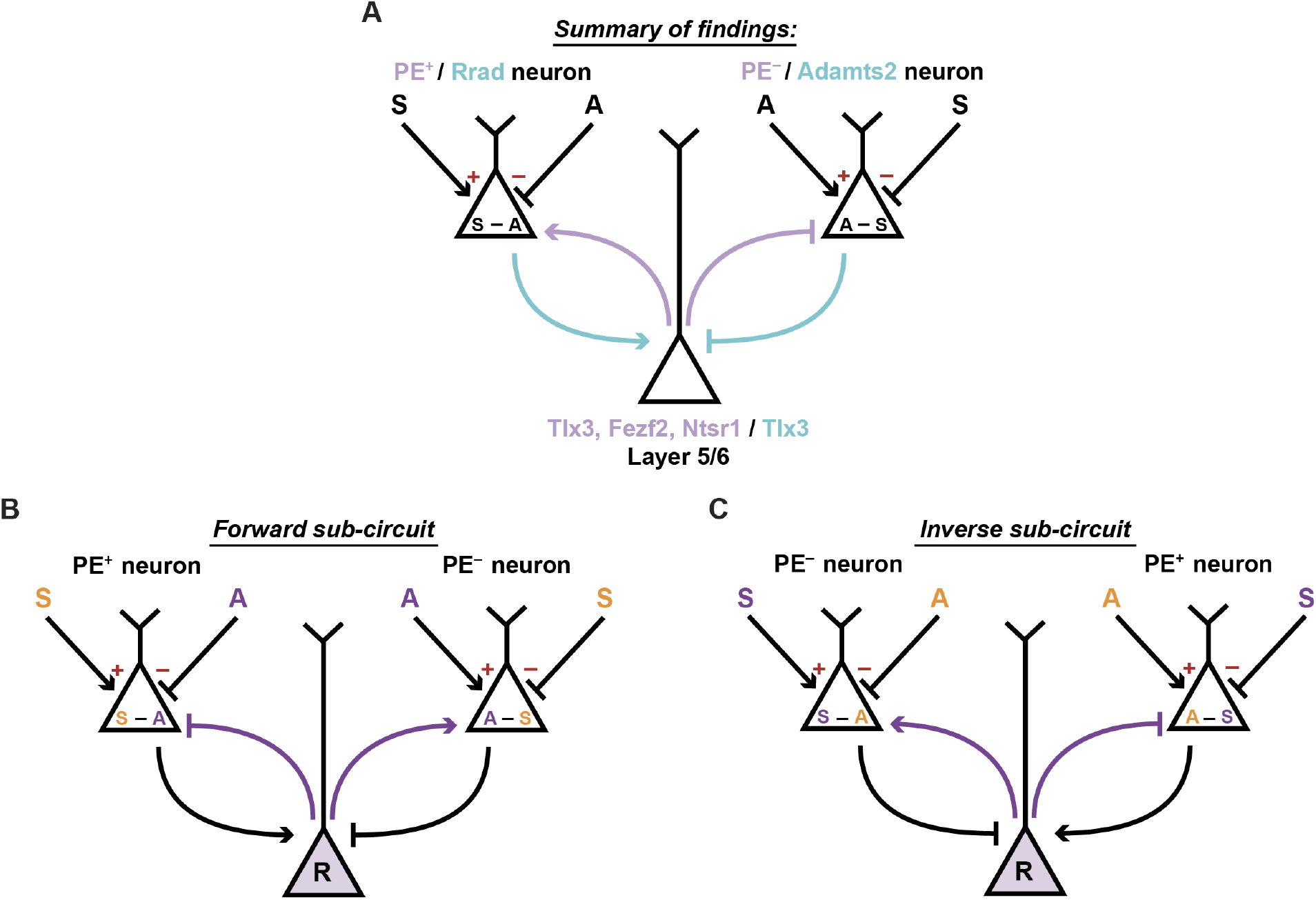
Sub-circuits for reciprocal predictive processing in a single cortical area. **(A)** Schematic of the functional influence patterns between cell types of deep and superficial cortical layers. Purple color marks functional connectivity from genetically identified cell types of deep layers on functionally identified prediction error types of layer 2/3. Blue color marks functional connectivity from molecularly identified cell types of layer 2/3 onto Tlx3 neurons of layer 5. **(B)** Schematic of the first (forward) sub-circuit. The action (A) input functions as a prediction and the sensory input (S) as a teaching signal. The internal representation neuron represents an estimate of the sensory input. **(C)** Schematic of the second (inverse) sub-circuit. The action (A) input functions as a teaching signal, and the sensory input (S) as a prediction. The internal representation neuron represents an estimate of the action.

## Key Resource Table

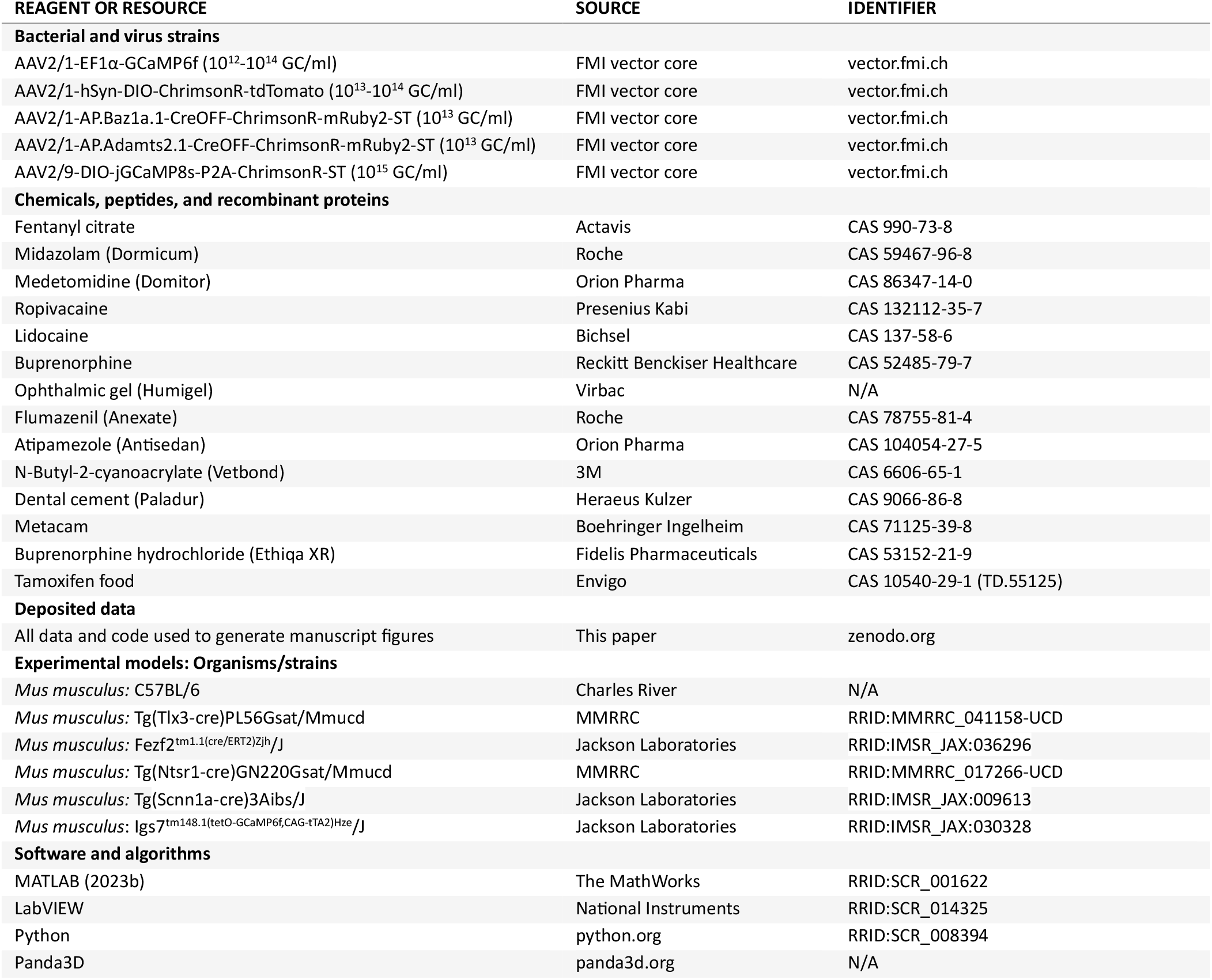

**Table S1.**
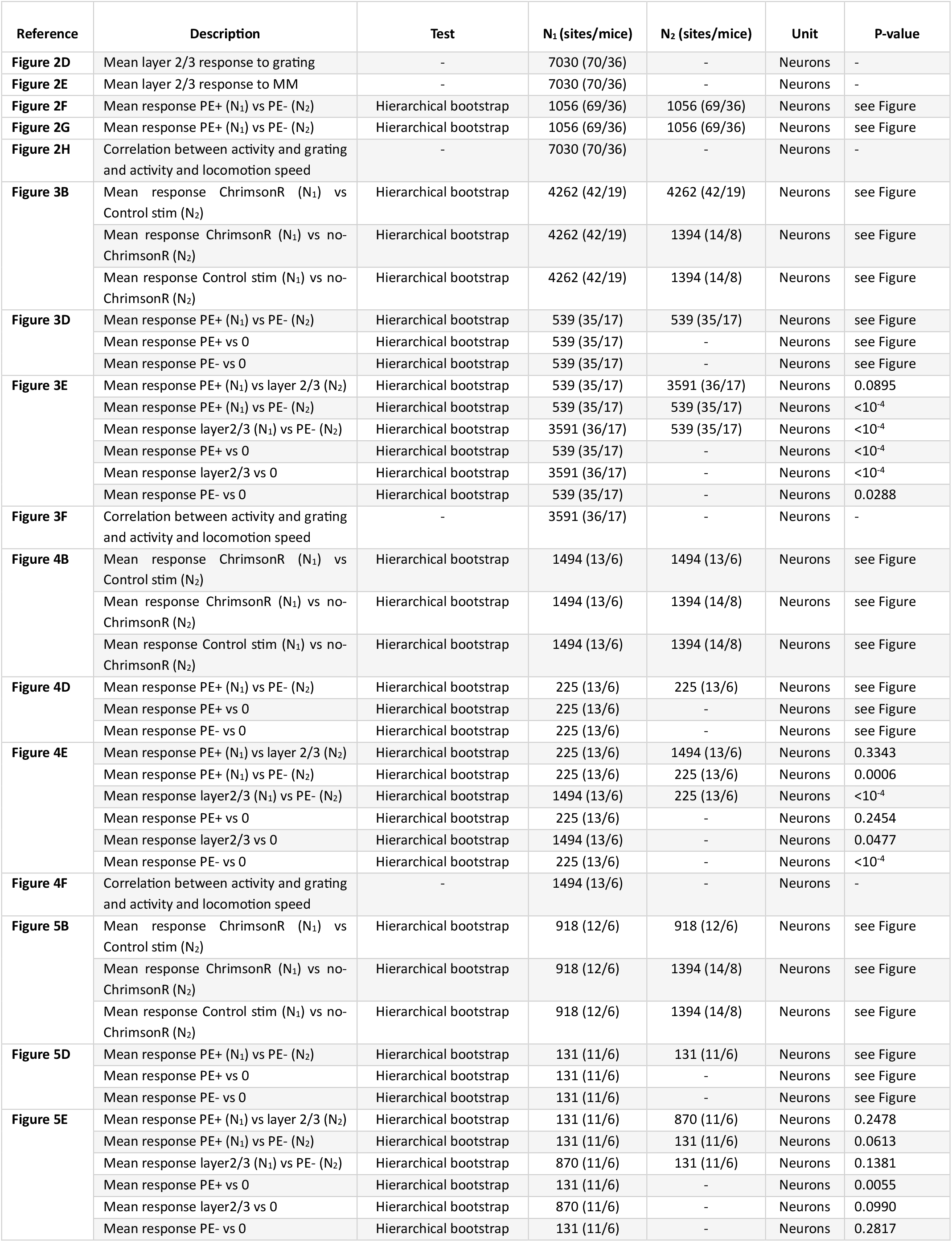
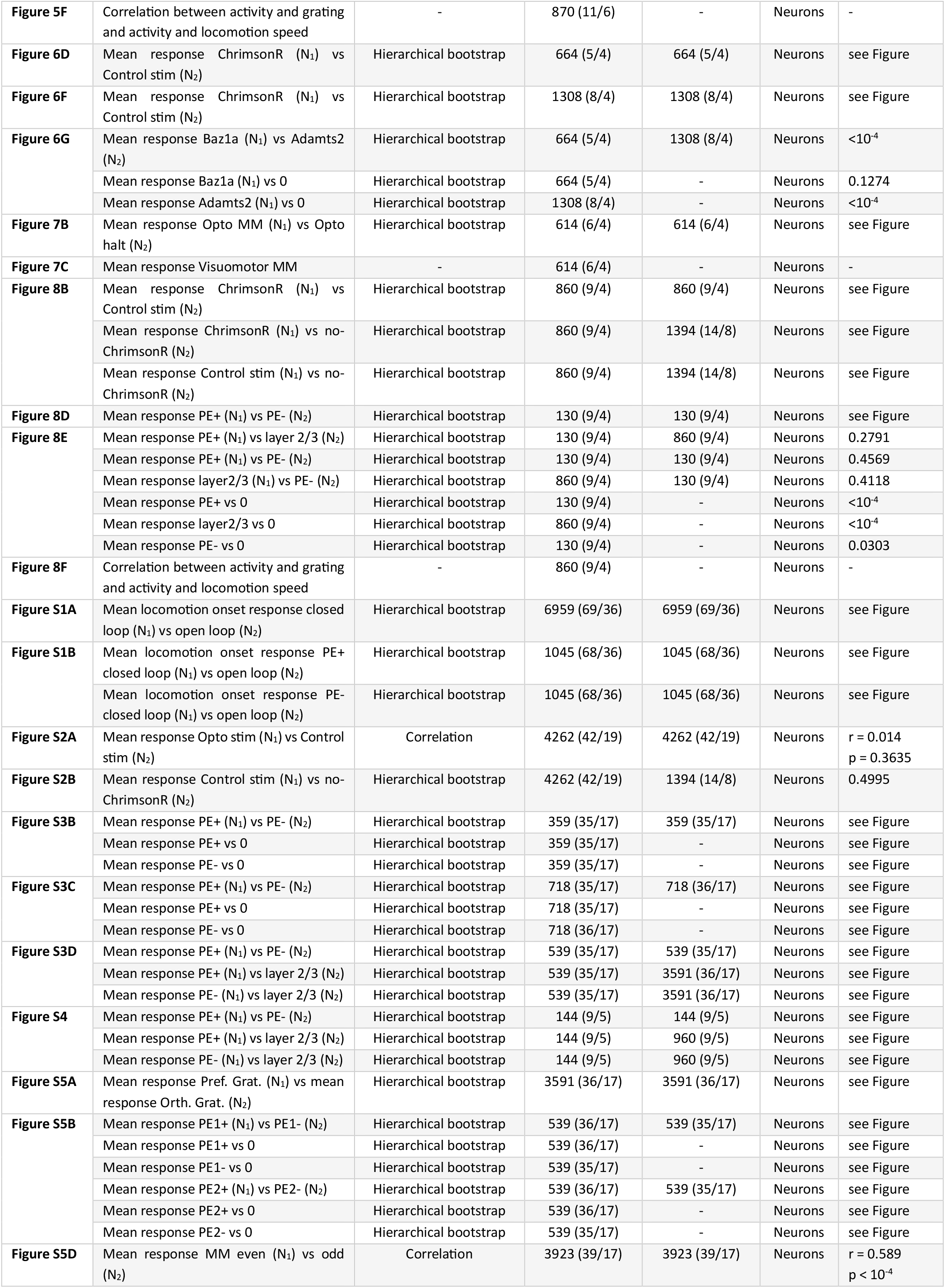
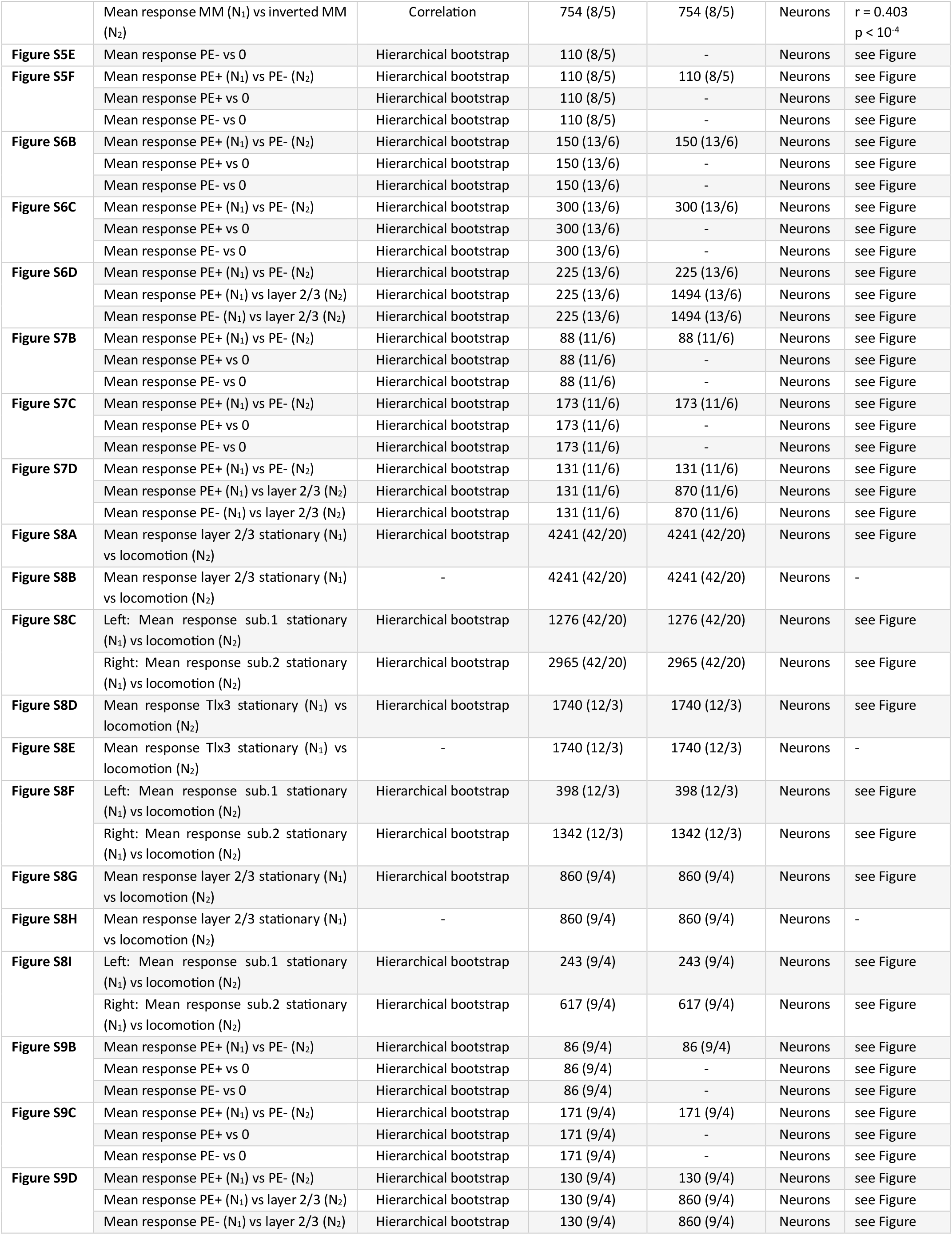
Statistics. All information on statistical tests used in the manuscript are shown in Table S1. We used hierarchical bootstrap or a correlation coefficient for all comparisons.

**Table S2.** Number of mice per experiment.

| Number of mice | Genotype | Vector | Figures |
| --- | --- | --- | --- |
| 10 | Tlx3-Cre | AAV2/1-EF1α-GCaMP6f,<br>AAV2/1-hSyn-DIO-ChrimsonR-tdTomato | Figures 2, 3B-3F, S1, S2, S3B-S3D, S5A-S5D, S8A-C |
| 5 | Tlx3-Cre | AAV2/1-EF1α-GCaMP6f,<br>AAV2/1-hSyn-DIO-ChrimsonR-tdTomato | Figures 2, 3B-3F, S1, S2, S3B-S3D, S5, S8A-S8C |
| 2 | Tlx3-Cre | AAV2/1-EF1α-GCaMP6f,<br>AAV2/1-hSyn-DIO-ChrimsonR-tdTomato | Figures 3B-3F, 7B-7D, S2, S3B-S3D, S5A-S5D, S8A-S8C |
| 1 | Tlx3-Cre | AAV2/1-EF1α-GCaMP6f,<br>AAV2/1-hSyn-DIO-ChrimsonR-tdTomato | Figures 3B-3C, 7B-7D, S2, S8A-S8C |
| 1 | Tlx3-Cre | AAV2/1-EF1α-GCaMP6f,<br>AAV2/1-hSyn-DIO-ChrimsonR-tdTomato | Figures 7B-7D, S8A-C |
| 1 | Tlx3-Cre | AAV2/1-EF1α-GCaMP6f,<br>AAV2/1-hSyn-DIO-ChrimsonR-tdTomato | Figures 3B-3C, S2, S8A-C |
| 3 | Tlx3-Cre | AAV2/9-DIO-jGCaMP8s-P2A-ChrimsonR-ST | Figure S8D-S8F |
| 6 | Fezf2-CreERT2 | AAV2/1-EF1α-GCaMP6f,<br>AAV2/1-hSyn-DIO-ChrimsonR-tdTomato | Figures 2, 4B-4F, S1, S6B-S6D |
| 6 | Ntsr1-Cre | AAV2/1-EF1α-GCaMP6f,<br>AAV2/1-hSyn-DIO-ChrimsonR-tdTomato | Figures 2, 5B-5F, S1, S7B-S7D |
| 4 | Scnn1a-Cre | AAV2/1-EF1α-GCaMP6f,<br>AAV2/1-hSyn-DIO-ChrimsonR-tdTomato | Figures 2, 8B-8F, S1, S8G-S8I, S9B-S9D |
| 5 | C57BL/6 | AAV2/1-EF1α-GCaMP6f | Figures 2, 3B, 4B, 5B, 8B, S1, S2B, S4 |
| 3 | C57BL/6 | AAV2/1-EF1α-GCaMP6f | Figures 3B, 4B, 5B, 8B, S2B |
| 4 | Tlx3-Cre x Ai148 | AAV2/1-AP.Baz1a.1-CreOFF-ChrimsonR-mRuby2-ST | Figures 6D, 6G |
| 4 | Tlx3-Cre x Ai148 | AAV2/1-AP.Adamts2.1-CreOFF-ChrimsonR-mRuby2-ST | Figure 6F, 6G |

